# Larval thermal tolerances and future dispersal potential of the subtropical-to-temperate sea urchin *Tripneustes australiae*

**DOI:** 10.64898/2026.10.13.758839

**Authors:** Emily McLaren, Paulina Selvakumaraswamy, Brigitte Sommer, Maria Byrne

## Abstract

The distribution and dispersal of marine ectotherms is mediated by their thermal tolerances, and as the oceans warm species distributions are shifting. Developmental thermal tolerances of marine invertebrates with a pelagic life-history phase can be used to predict the potential for local extinctions and range extensions. In southeast Australia, *Tripneustes australiae* is an ecologically important sea urchin species as it undergoes periodic outbreaks and can transform ecosystems through intense herbivory. We investigated the embryonic and early larval developmental tolerances of *T. australiae* with respect to its current distribution, to predict current and future dispersal capacity. The optimal thermal window for survival, normal development and developmental progression of *T. australiae* ranged between 17 and 23°C, with probability of survival at these temperatures remaining above 85% throughout the experiment. This developmental thermal window overlaps with the current thermal regime experienced across the adult distribution. This suggests that *T. australiae* may have a similar thermal window in its adult and early larval life history stages. Further, *T. australiae* may have the potential to persist at its warm-range edge into the future and to disperse and range shift poleward in coming decades. Indeed, we show that the temperature at which larvae have a 50% probability of survival may extend poleward across the Bass Strait by 2100. The persistence of *T. australiae* in its mid-range, its potential to disperse poleward, and increased grazing pressure from periodic outbreaks, may impact temperate macroalgal habitats already at risk from warming and concentrated grazing by other sea urchin taxa.

## Introduction

Species distributions are mediated by their thermal tolerances (Sunday et al. 2011; Stuart-Smith et al. 2017). As the oceans warm, marine communities are reorganising and species are moving to track changes in their thermal environment (Pinsky et al. 2013; García Molinos et al. 2016; Poloczanska et al. 2016; Stuart-Smith et al. 2022; Fredston et al. 2025; Oliveira et al. 2026). This global phenomenon is accelerating in regions with strong continental boundary currents that transport propagules poleward (Vergés et al. 2014, 2019; Zarzyczny et al. 2023). Species with generalist traits, such as broad thermal tolerances, are more likely to undergo range expansions (Bates et al. 2014; Sunday et al. 2015; Byrne et al. 2022). Importantly, cool thermal tolerance limits can indicate a species’ ability to range extend poleward, and thermal minima can be plastic (Sandblom et al. 2016; Byrne et al. 2017).

Many marine invertebrates have a bi-phasic life history that comprises a dispersive pelagic larva and benthic adult (Strathmann 1993; McEdward and Miner 2001). The thermal tolerance of the larval stage can be used to predict species distributions, range extension capacity and local extinction risk (Byrne 2011b; Byrne et al. 2016; Collin et al. 2021; Byrne et al. 2022). As such, larval thermal tolerances have been the focus of many studies in the context of climate change (Byrne 2011b; Byrne and Przeslawski 2013; Gall 2021; Lang et al. 2023) as marine heatwaves and ocean warming become more prevalent (Tuckett et al. 2017; Edgar et al. 2023). Indeed, for marine ectotherms the pace of developmental progression is governed by temperature. Faster development in warmer conditions can reduce the pelagic phase and the ‘peril of the plankton’ which may increase survival of propagules (Vaughn and Allen 2010). While there tends to be a focus on warm tolerance, understanding cool-tolerance limits is essential to predict cool-edge range extensions (Bates et al. 2014; Sunday et al. 2015; Byrne et al. 2017, 2022; Stuart-Smith et al. 2017, 2022; Edgar et al. 2023). This is particularly important to understand for taxa such as sea urchins, which can have long planktonic durations and the ability to transform ecosystems through poleward range extension and concentrated herbivory (Ling 2008; Valentine and Edgar 2010).

Echinoids are important herbivores that contribute to the barrens - kelp forest habitat mosaic characteristic of nearshore temperate marine communities (Steneck 2020), including in southeast Australia (Jones and Andrew 1990; Underwood et al. 1991; Przeslawski et al. 2023). Southeast Australia is a hotspot for echinoid diversity, with species from diverse families occupying rocky reef, seagrass, and soft-sediment habitats (Byrne and O’Hara 2017; McLaren et al. 2026). However, this biodiverse region is also an ocean warming hotspot (Hobday and Pecl 2014), where the East Australian Current is driving rapid warming (Suthers et al. 2011) and facilitating poleward range extensions of many taxa (Ling 2008; Vergés et al. 2014, 2019).

The poleward range extension of the diadematid sea urchin *Centrostephanus rodgersii* from New South Wales (NSW) to Tasmania across the Bass Strait has transformed receiving seascapes from kelp forests to sea urchin barrens habitat (Ling 2008). This change was driven by the increasingly favourable thermal conditions for the poleward larval dispersal of *C. rodgersii* and post larval success (Byrne et al. 2022). In contrast, other sympatric echinoids with dispersive larvae that are common and abundant in southeast Australia, such as *Heliocidaris tuberculata* and *Tripneustes australiae*, have not range extended to Tasmania. For *H. tuberculata* this is due to intolerance of larvae to cooler higher latitude temperatures (Byrne et al. 2022). As such, understanding the cool thermal tolerances of echinoid taxa in southeast Australia is ecologically important.

*Tripneustes* sea urchins are globally important herbivores in seagrass and rocky reef habitats and are characterised by boom-bust population dynamics that can transform ecosystems (Uthicke et al. 2009; Valentine and Edgar 2010; Lawrence and Agatsuma 2020; Moreira- Saporiti et al. 2023; McLaren et al. 2024; Byrne et al. 2026). In southeast Australia, the local *Tripneustes* is *T. australiae,* commonly referred to as the ‘Lamington’ sea urchin (Bronstein et al. 2017, 2019; McLaren et al. 2023, 2025). *Tripneustes australiae* is an ecologically important species known to have population outbreaks that reduce macroalgae cover in subtropical and temperate habitats (Valentine and Edgar 2010; Byrne et al. 2024,2026; McLaren et al. 2024).

We investigated the developmental thermal tolerances of *T. australiae* with respect to their realised thermal niche as determined from its adult distribution in Australia (Figure 1 and see McLaren et al. 2025). This species’ adult distribution in Australia ranges from its northern warm edge around Byron Bay, NSW (28°37’S) to its southern cool limit in Narooma (36° 15’S). Based on the climatology where adult *T. australiae* have been observed at their range edges, this species has an estimated realised thermal niche of 14.2 - 26.6°C (Figure 1; McLaren et al. 2025). While adult distribution provides information on where the benthic stage has been successful, the larval stage may have a different, potentially broader, thermal tolerance (Lamare et al. 2018; Collin et al. 2021). We conducted an experiment to understand the developmental thermal tolerance of *T. australiae* and the influence of temperature on progression through developmental stages. We predicted that given the adult distribution, the lower and upper thermal tolerance of development would approximate 14°C and 26°C, respectively. Considering the dynamic warming of southeast Australia, we then used the warm and cool developmental thermal tolerances from our experiment to determine the current and future dispersal potential of *T. australiae* in southeast Australia.

**Figure 1.**
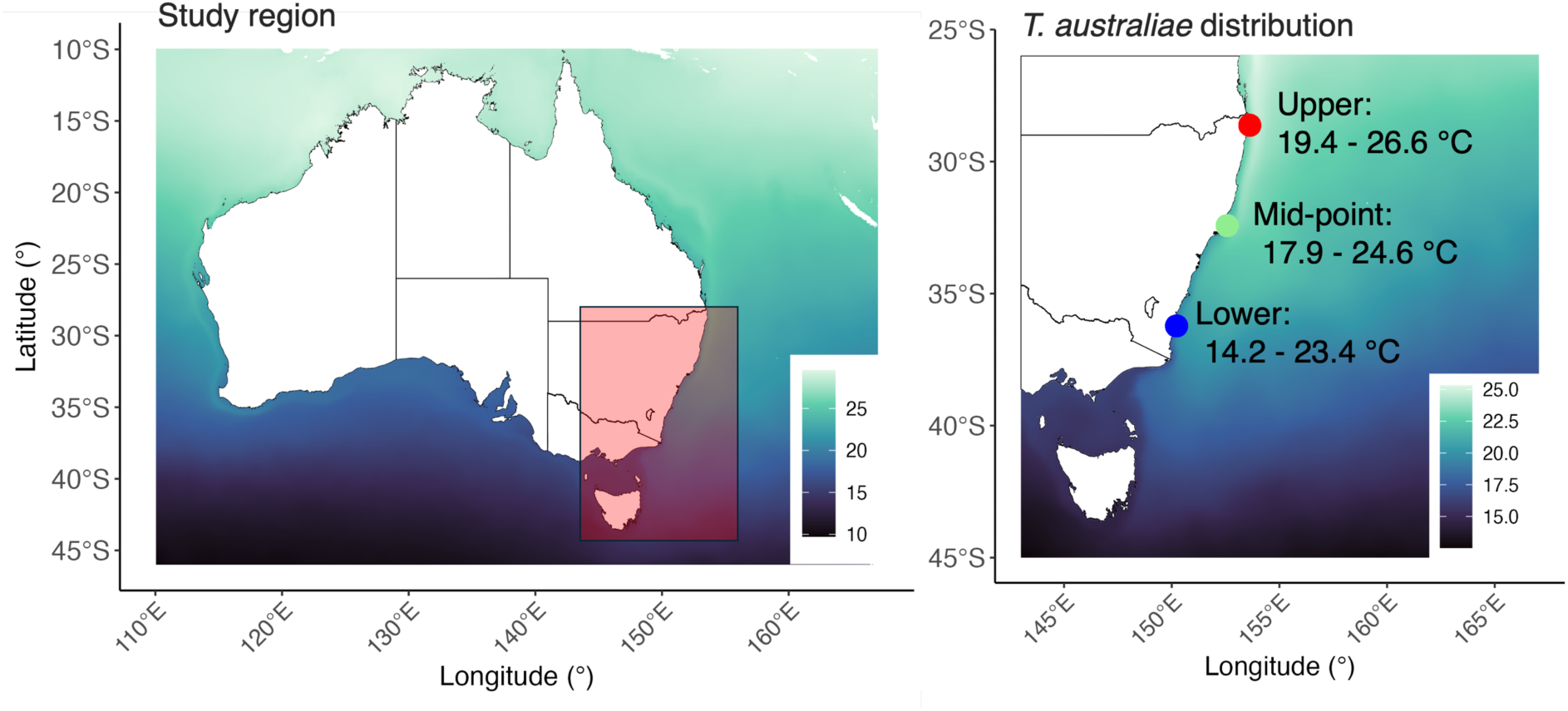
Study region in southeast Australia (left) and the adult distribution of *Tripneustes australiae* in Australia (right). The upper, lower, and mid-range points are Julian Rocks (28°37′30″ S), Forster (36°13′12″ S), and Montague Island (32°25′30″ S) in New South Wales. Temperature ranges represent the average highest and lowest temperature records for each year in the periods 2000-2010 and 2010-2020 (°C) (Assis et al. 2024). Background colours represent the mean SST from 2000 to 2020 extracted from Bio- Oracle SST (Assis et al. 2024).

## Methods

### Specimen collection, spawning and larval rearing conditions

*Tripneustes australiae* (n ∼ 30) were collected from Port Stephens, New South Wales (32.715° S, 152.182° E) (2-5m depth) in June 2023 (Test diameter: 63 mm – 83mm). At the time of collection, the ambient sea water temperature was 18.4°C (±0.4) (Beachwatch NSW 2024). Animals were transported to the University of Sydney and maintained at 20°C in aquaria and fed *Sargassum* sp. *ad libitum* for ∼30 days until spawning.

Spawning was induced by gentle manual manipulation (oral side up to oral side down). Gametes were collected on release from the gonopore with sperm placed into 1.2 ml tubes and stored at 4°C until fertilisation and eggs transferred into beakers (1000 ml) filled with 1μm filtered sea water (FSW) (20°C). Gamete quality was checked microscopically to ensure sperm motility and that the eggs had a regular shape and lacked germinal vesicles. Two independent fertilisations were undertaken. For each fertilisation we pooled sperm from 3 males (M) and eggs from 2-3 females (F) (Fertilisation 1: 3M, 3F; Fertilisation 2: 3M, 2F). Fertilisation success (90%) was checked microscopically for the presence of the fertilisation envelope in a random subsample of 100 embryos. The cultures (15 embryos/larvae ml^-1^) were established using routine methods (Byrne et al. 2022). From these cultures cleavage stage embryos (2-4 cell stage) were transferred into the vials and placed in the experimental temperatures (see below). Vials with embryos from each independent fertilisation were randomly interspersed throughout the experimental treatments. Every 48 hours, half of the FSW in the vials was replaced with new FSW at the experimental temperature. As *T. australiae* is known to have a long facultative feeding period (∼ 8 days) at ambient temperature 19-20°C (Byrne et al. 2008a, b), the larvae were not fed for the duration of the experimental period. Salinity and dissolved oxygen (DO) were monitored regularly (WTW Multi 3420; Thermo Fisher Scientific Eutech DO 6+) and remained at ∼35 PSU and >90% DO.

### Experimental design

The effect of temperature and time (days post fertilization, DPF) on survival, normal development and developmental stage was assessed using five temperature treatments (14°C, 17°C, 20°C - control, 23°C and 26°C), at seven time points (1-, 2-, 3-, 5- and 7-days post fertilisation) from cleaving embryos and blastula to echinopluteus larva. Temperature treatments were chosen to reflect the climatology where adult *T. australiae* have been observed at their range edges (14.2 - 26.6°C) (Figure 1; McLaren et al. 2025). Treatment temperatures were established using water baths and each treatment temperature had four independent replicates (vials) for each time point (4 vials x 5 temperatures x 5 time points; n = 120). At each time point four vials were removed for scoring and then discarded to avoid repeated measures and effects of handling. Water bath temperature was measured at each scoring time point (n = 7) to ensure they remained stable (13.9°C ± 0.16; 16.96°C ± 0.10; 19.98°C ± 0.33; 22.91°C ± 0.73; 25. 89°C ± 0.04).

### Larval scoring and survival

Prior to scoring, the embryos/larvae were dispersed in the vial by gentle inversion, and a random sample of the first 30 embryos/larvae were scored microscopically for survival (dead or alive), condition (normal or abnormal embryos/larvae) and developmental stage. Across the treatments and time points, development was scored according to stage as cleaving (2-cell – morula), blastulae (unhatched and hatched), gastrulae (early through late), prism or pluteus (2- and 4-arm echinoplutei) (Figure 2). Survival was determined by the presence of motile embryos/larvae. Dead embryos/larvae were immotile or disintegrating. Abnormal embryos and larvae had a range of phenotypes including specimens with an incomplete archenteron or abnormal larval profile (Figure 2). Scoring was not performed blind to treatment. Due to high mortality and likely dissolution (disappearance) of dead larvae by day 9, the data for this time point were not used in the assessment of the percentage of normal development or for survival analysis. However, the day 9 data were used to construct thermal performance curves (TPCs) to model 50% mortality.

**Figure 2.**
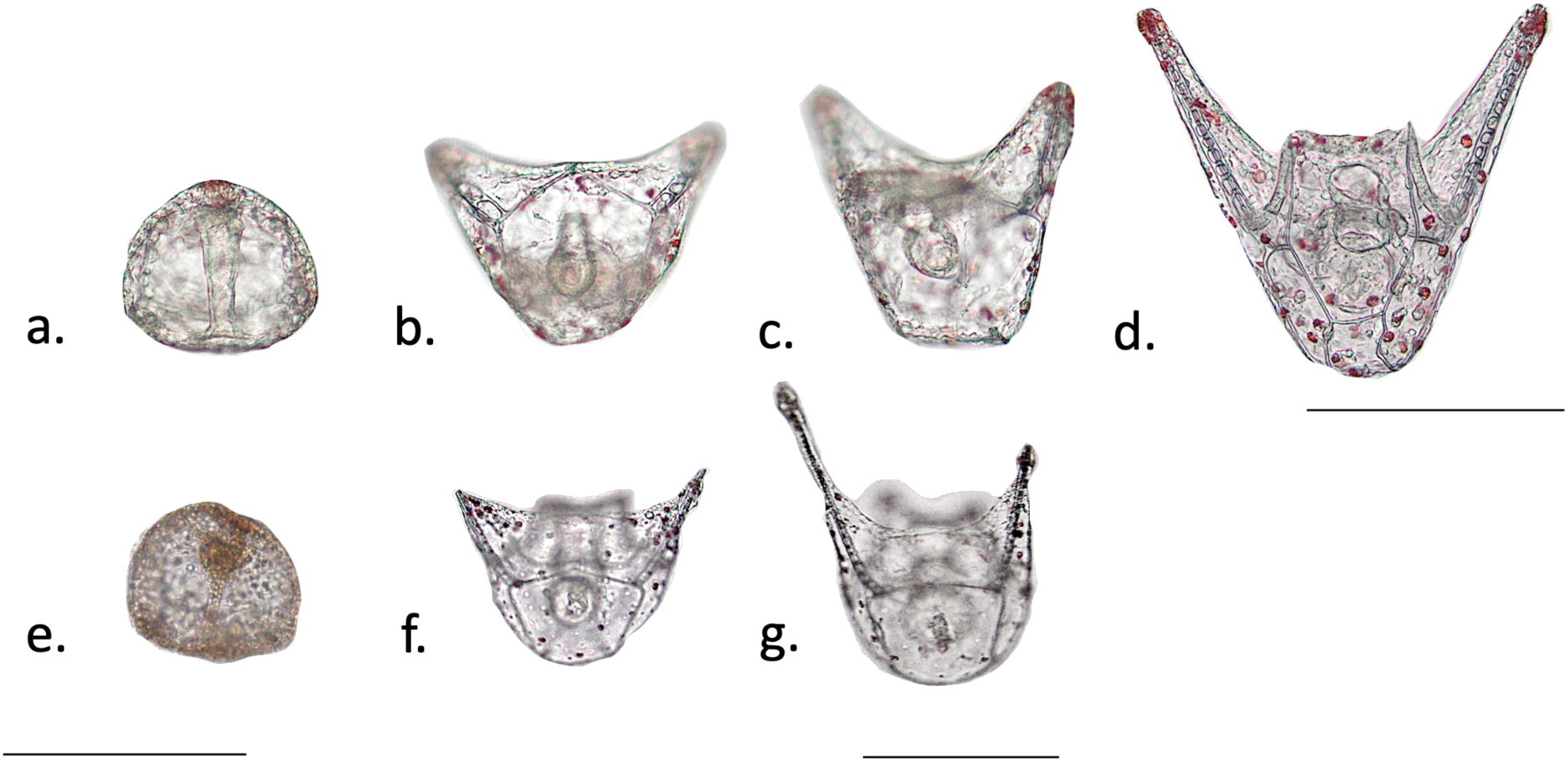
Normal embryo/larval profiles of *T. australiae* for gastrula (a.), prisms (b. & c.) and 2-arm plutei (d.). Abnormal embryo/larval profiles of T*. australiae* for gastrula (e.), late prisms/2-arm plutei (f.) and later stage 2-arm plutei (g.). Scale bars: a-g = 500µm, e = 200µm.

### Statistical analysis

#### Survival

The proportion of live or normal larvae was determined as the number of larvae alive (n = 30) in each replicate (n = 4) for each day post fertilisation (DPF) (n = 5) and temperature (n= 5).

To determine the effect of temperature and DPF on the probability of survival and normal embryonic / larval development, we used two separate Generalised Linear Mixed Models (GLMM) with a beta-binomial distribution in the *glmmTMB* package (Brooks et al. 2017). DPF (continuous 1-7) and temperature (ordered categorical) were included as fixed effects and the random effect of replicate was included to account for any variation among the vials (model: alive, dead or normal, abnormal ∼ temperature * DPF + (1| replicate)). Initial models were built with an interaction term between DPF and temperature to explore an interactive effect of these fixed effects. We tested model assumptions and performed model checks for overdispersion and outliers using the *DHARMa* package (v. 0.4.7) (Hartig 2026). Model significance was tested with Type II Wald Chi-square Tests using the *Anova* function in the car package (Fox and Weisberg 2019). Post-hoc comparisons between temperatures and timepoints were performed using the *emmeans* package (Lenth 2024).

#### Thermal performance curves (TPCs)

Data on percentage survival at days 5, 7, and 9 were analysed using separate general additive models (GAM) in the *mgcv* package (Wood 2017) to create thermal performance curves (TPCs) and determine the upper and lower lethal temperature for 50% mortality (LLT_50_ and ULT_50_) on these days. The percentage of live larvae was modelled as a function of a spline smoothed term for temperature with *k* (basis dimension) set to 4 for day 5 and 9, and 3 for day 7. Both GAMs were fit using the Restricted Maximum Likelihood (REML) method. Model fit was checked using the *gam.check* function and visually inspecting the Q–Q plots, and the spread of residuals was assessed using residuals versus fitted values. TPCs with 95% confidence intervals were then constructed from the predicted values of the GAMs and LLT_50_ and ULT_50_ were determined from these predicted values.

#### Developmental thermal tolerance

To examine how developmental stage progression differed across the temperature treatments and timepoints we used a multivariate approach which allowed developmental stage to be treated as multivariate response data. We did so to test the multivariate hypothesis that the number of larvae in respective developmental stages would vary across the temperatures and timepoints, and to test for an interactive effect between DPF and temperature. To do so we fit multiple Generalised Linear Models (GLMs) simultaneously using the *mvabund* package (v. 4.2.8) (Wang et al. 2026). The model explored the number of larvae at each developmental stage as a function of temperature and day with a negative binomial distribution and a log-link function (model = number of larvae at developmental stage ∼ temperature*DPF). For multivariate count data, Warton et al. (2005) show that the negative binomial distribution is the best model structure and so this is the default implementation in *mvabund*. More so, the *mvabund* function estimates an overdispersion parameter for each model variable using the maximum likelihood estimations, accounting for any overdispersion in the model. Assumption and model checks were undertaken by visually inspecting the spread of residuals using a residuals versus fitted values plot. The *anova* function in the *mvabund* package was used to test the significance of this model which performs a likelihood ratio test and reports resampled p-values. To examine the hypothesis further and understand which developmental stages are likely to be at each timepoint at different temperatures we used a series of univariate GLMs within the package *mvabund* (Wang et al. 2026). Each of these univariate GLMS had the same model structure as above, but was performed for each developmental stage (embryos, blastula, prism or pluteus). The adjusted re-sampling method in the *anova* function was used to calculate p-values, which accounts for any correlation between the developmental stages in the vials. All statistical analyses were performed in R (R Core Team 2025).

### Current and future potential larval dispersal

To determine current and future dispersal potential of *T. australiae* in southeast Australia, the upper and lower lethal temperatures for 50% mortality (ULT_50_:23.7°C; LLT_50_: 15.82°C) determined from the thermal performance curves were aligned with remotely sensed sea surface temperature (SST, °C) data extracted from the marine data repository Bio-Oracle (Assis et al. 2024). We extracted the 2000–2010 and 2010–2020 summary data sets from the Bio-- Oracle SST records (Assis et al. 2024) and took the average of the two average temperatures for each decade to determine overall average for the 2000–2020 period. To determine future dispersal potential, we extracted the 2090–2100 period under the IPCC concentration pathway SSP2-4.5. This pathway represents a ‘middle of the road’ scenario, assuming a conservative 2.7°C rise in global temperature by 2100 (IPCC 2023). The latitude and longitude at which the ULT_50_ (23.7°C) and LLT_50_ (15.82°C) currently occurs and will occur in 2100 was then mapped to determine where current and future dispersal limits occur in our study region.

## Results

### Survival

There was a significant interaction between temperature and day on the probability of live *T. australiae* embryos/larvae (χ^2^ = 17.184, df = 4, p = <0.001). For larvae reared at 14°C, the probability of survival was above 85% until day 3, when probability of survival decreased to 66%. At this temperature, by day 5 larvae had a 10% chance of survival and on day 7 there was 100% mortality (Figure 3). This indicates the 14°C temperature treatment is lethal to *T. australiae* larvae. Further, for the 14°C rearing temperature all days were significantly different to each other (p < 0.001, Table S1) (Figure 3). There was no significant variation in survival through time (p>0.05, Table S1) for the 17 °C, 20°C, and 23 °C treatments, with the probability of survival remaining above 85% for the entire sampling period. Larvae reared at the warmest temperature 26°C had a 90% probability of survival until day 5. Thereafter, these larvae had significantly lower survival, decreasing to 70% on day 5 (p < 0.05) (Figure 3), and to 30% probability of survival on day 7 (p < 0.001, Table S1).

**Figure 3.**
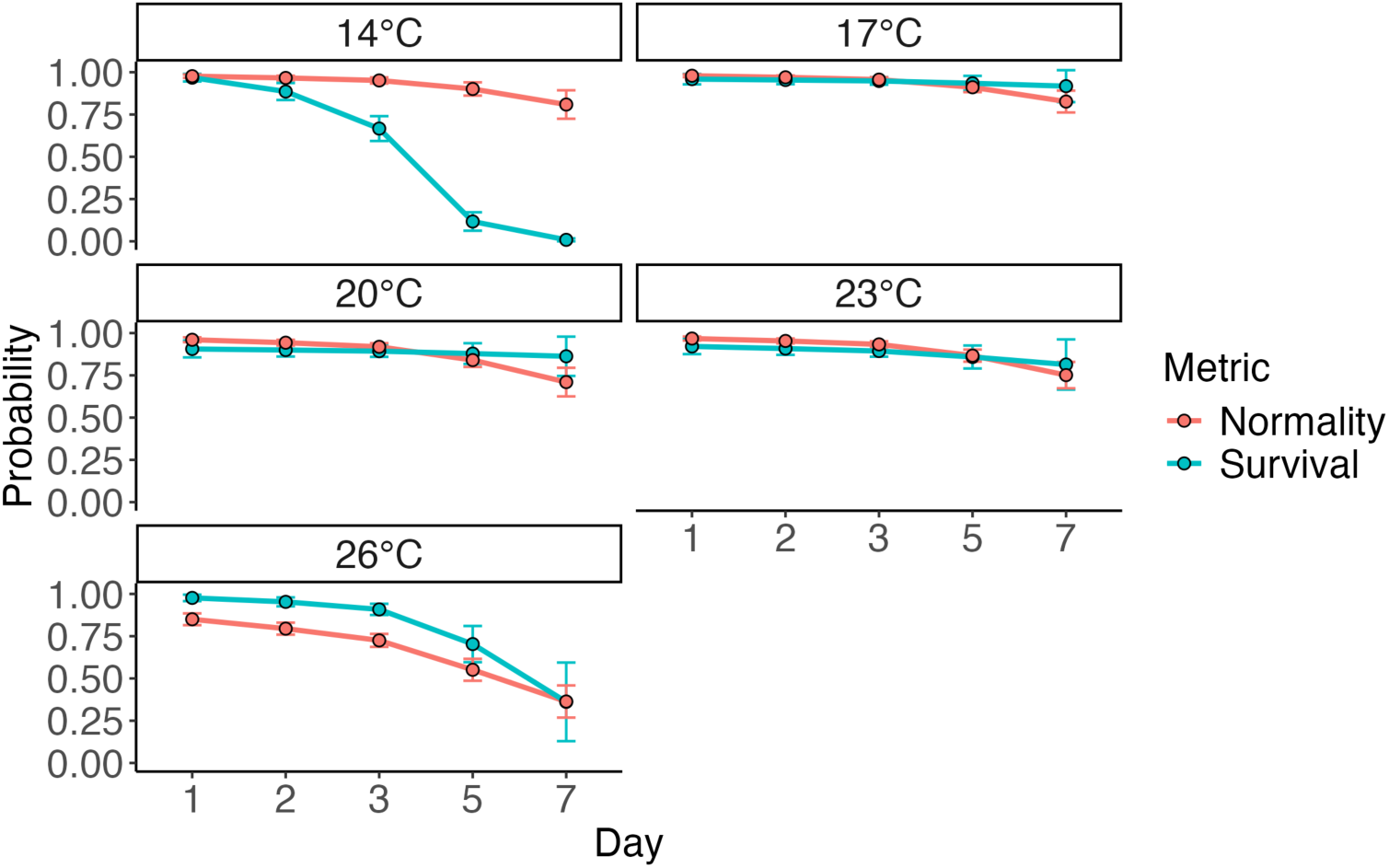
Model predicted probability of survival and normal development (±SEM) of *Tripneustes australiae* for the five time points (1-7 days post fertilisation) across five rearing temperatures (14, 17, 20, 23, & 26°C).

### Normal development

Contrary to survival, there was no significant interaction between DPF and temperature on the probability of normal development (Figure 3) (χ^2^ = 5.549, df = 4, p >0.05). Instead, the proportion of embryos/larvae with a normal profile varied significantly with days post fertilisation (χ^2^ =16.985, df = 1, p <0.001) and temperature (χ^2^ =48.382, df = 4, p <0.001). The probability of normal development decreased from 96% on day 1 to 70% on day 7, and all days differed from each other (averaged across temperature treatments, p<0.001, Table S2). The probability of normal development was 96% at 14 °C, this decreased to 74% at 26°C (averaged across DPF). The temperature treatments 14°C, 17°C and 23°C were all significantly different to the 26°C treatment (p<0.001, Table S3). At the coolest rearing temperature 14°C, all larvae were dead by day 7, however until that timepoint the probability of normal development was 96%. As such, the embryos in 14°C may have reached the gastrula stage and arrested development due to the cold (see below). Therefore, successful normal development for this species seems to occur between 17 and 23°C.

### Thermal performance curves

There was a significant relationship between larval survival and temperature on day 5 (F = 5.5, edf = 2.231, p = 0.0143). The lower lethal temperature limit (LLT_50_) was 14°C (95% CI: 32.6 - 72.8%), but only 43% of the variation was explained by the model, suggesting a weak relationship between temperature and survival at this time point. There was a significant relationship (F = 16.25, edf = 1.958, p<0.001) between temperature and the survival of *T. australiae* at day 7, with 66.9% of the variation explained by temperature. Between days 5 and 7 the predicted temperature at which 50% (day 5 95% CI: 32.6 - 72.8 %; day 7 95% CI: 65 – 35 %) of the larvae died increased from14°C to 15.94°C (Figure 4). On day 9, there was a significant relationship between temperature and survival (F = 58.92, edf = 2.915, p<0.001; 92.1% variation explained), and the LLT_50_ (day 9 95% CI: 41.9 - 60.1%) decreased to 15.81°C. Conspicuous mortality was observed in the upper temperature treatments on day 9, with 50% mortality (ULT_50_) (95% CI: 41.8- 61.5)% at 23.7°C. The thermal performance curves for survival of *T. australiae* narrowed through time decreasing from a lower limit of 15.94°C and no upper limit on day 7, to 17 - 23°C on day 9 (Figure 4).

**Figure 4.**
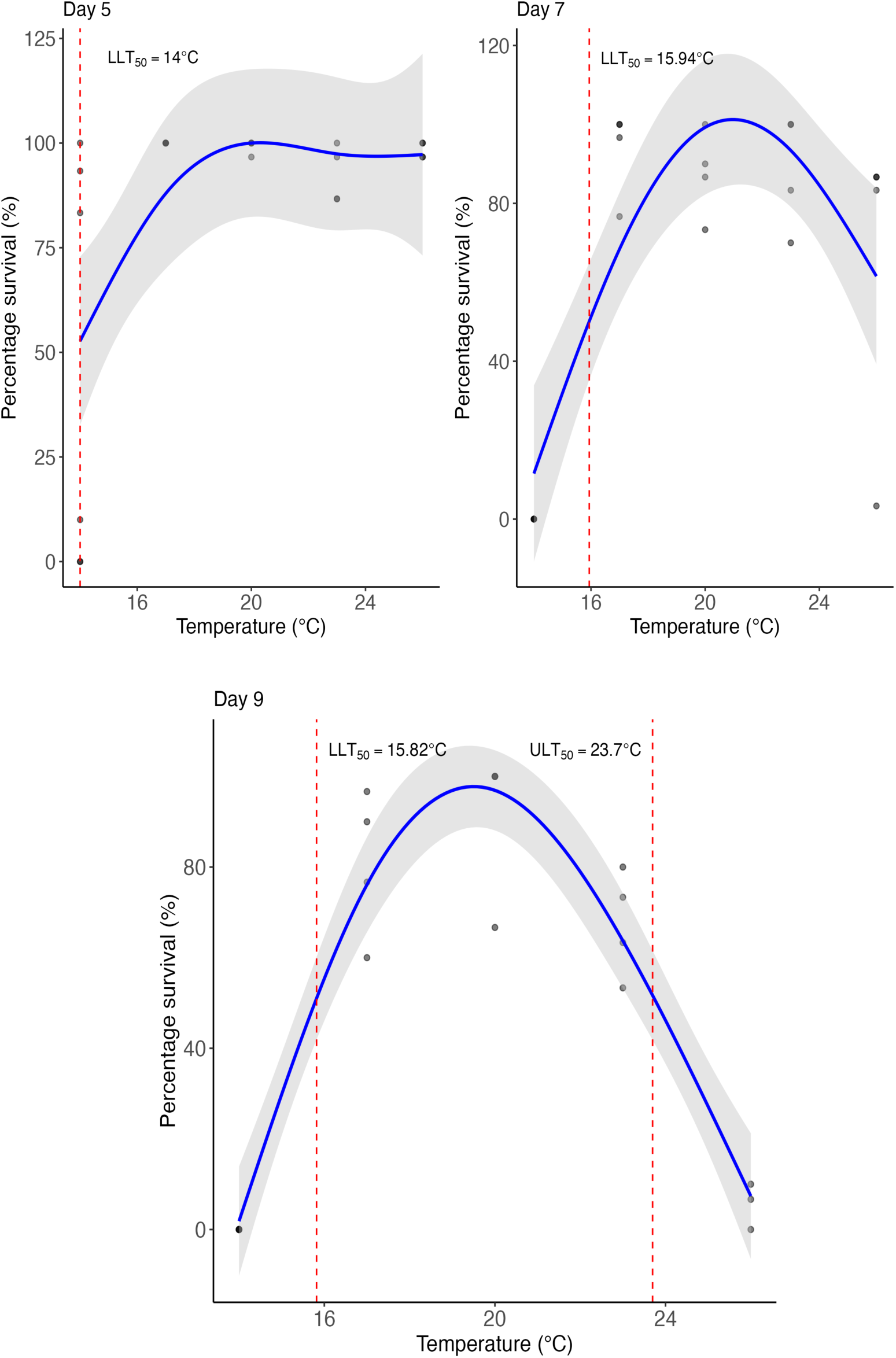
Thermal performance curves (95% CI) for *Tripneustes australiae* larvae constructed from general additive models (GAMs), for days 5, 7 and 9 post-fertilisations. The upper and lower LT50 temperatures (at which 50% survival occurs) are indicated by the red dashed line. The shaded band indicate the 95% confidence intervals.

### Developmental stage progression

With respect to larval stage there was a significant interaction between temperature and time (LRT = 343.2, res df = 85, df = 20, p < 0.001). Embryos/larvae were at different developmental stages across time points depending on rearing temperature, with embryos reared at warmer temperatures progressing through developmental stages faster (Figure 5).

**Figure 5.**
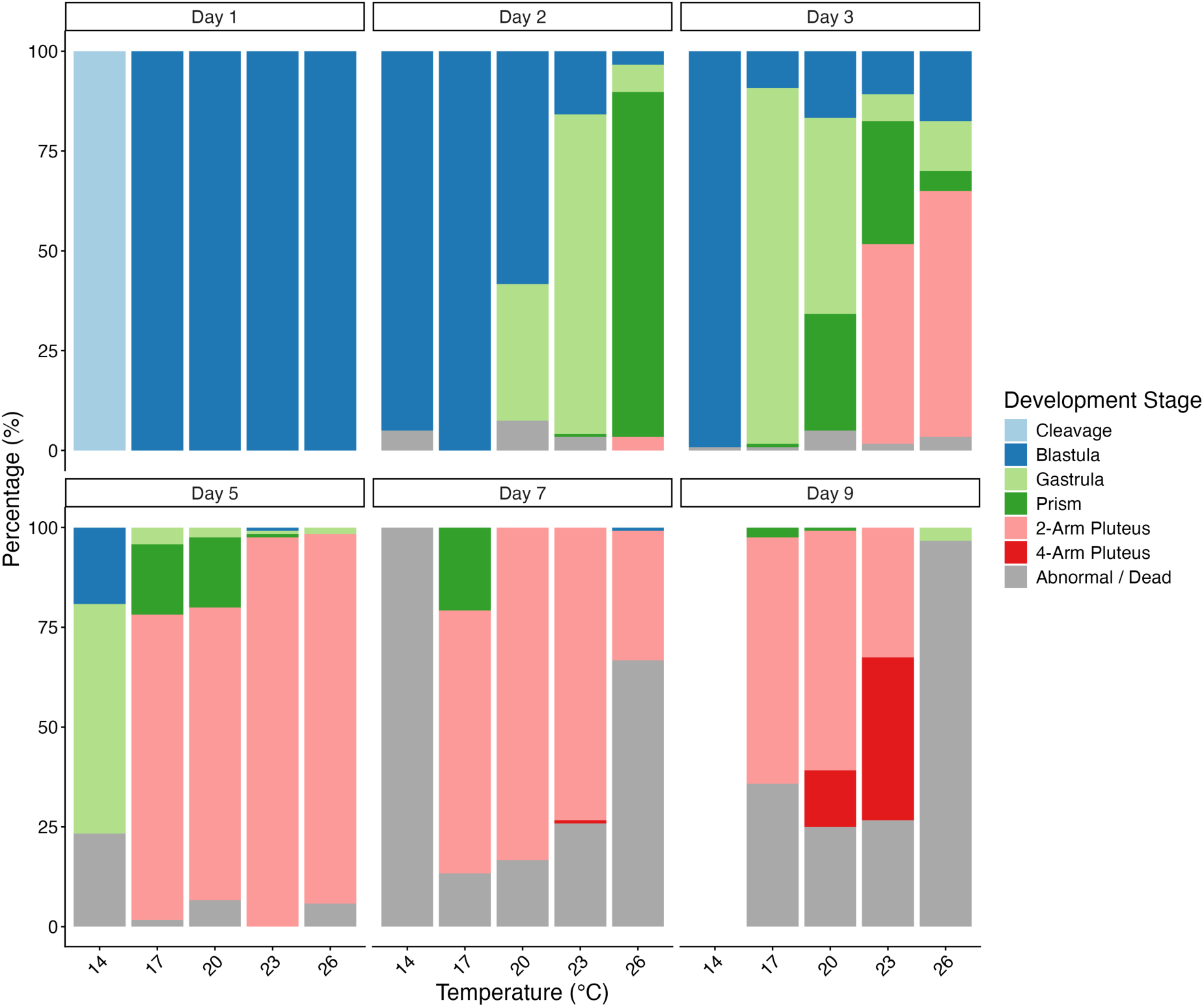
The percentage of different developmental stages of *Tripneustes australiae* for six time points post-fertilisation across five rearing temperatures (n = 30). Means and standard error for the respective treatments and timepoints can be found in Table S4.

The number of cleavage stage embryos present depended on the sampling day, with no influence of temperature (LRT = 40.21, p = 0.001) and they were only observed in the 14°C group at 24 hours. With respect to number of blastulae, gastrulae, prisms and plutei, there was a significant interaction between day and temperature (blastula: LRT = 118.9, p = 0.001; gastrula: LRT = 84.98, p = 0.001; prisms: LRT = 48.9, p= 0.002; 2-arm plutei: LRT = 62.6, p = 0.001). There was no significant effect of day, temperature or their interaction on the number of 4-arm plutei (day*temperature: LRT = 0.05, p =0.291; day: LRT = 19,35, p = 007; temperature: LRT = 9.4, p = 0.101). On day 1, temperature groups between 17°C and 26°C were blastulae, and on day 3, only the 14°C degree group were primarily blastulae (Figure 5, Table S4). Gastrulae first appeared on day 2 in the 20-26°C temperature treatments with the highest proportion of this stage in the 23°C group (Figure 5, Table S4). In the warmest treatment (26°C) on day 2, the cultures were at the prism stage, having completed gastrulation by 48 hours post-fertilisation (Figure 5). In comparison, controls were primarily blastulae on day 2, and some had gastrulated (Figure 5). By day 3, the 23°C and 26°C treatments had 50% plutei and by day 5 the 17-26°C treatments were dominated by plutei (Figure 5, Table S4). By day 9 4-armed plutei were present in the 20 and 23°C treatments The 14°C group never achieved the pluteus stage and 26°C group had near total mortality at day 9.

### Current and future potential larval dispersal

In southeast Australia, the current dispersal potential of *T. australiae* with respect to the average sea surface temperature and the developmental lower thermal limit (LLT_50_) seems to be mediated by the East Australian Current (EAC) in the Bass Straight (Figure 6). The warm and cool range limits where 50% survival through early development is possible, are currently at 25.975°S and 42.275°S, respectively. In 2100, the range edges shift poleward to 26.425 °S at the cool range edge and to 43.575 °S at the warm range edge. As 1° latitude corresponds to ∼111km distance, the warm and cool range edges will therefore shift poleward by approximately 50 km and 144.5 km, respectively. Although currently, propagules may disperse in the EAC across the Bass Straight, the average temperatures of nearshore Tasmania and islands in the Bass Straight are not suitable for successful development (Figure 6). However, in 2100 the nearshore habitat in northeastern Tasmania and the islands in Bass Straight may become thermally suitable for the successful development of *T. australiae* to the echinopluteus stage (Figure 6). Further, the latitude for the upper thermal limit (ULT_50_) of development may move poleward by 2100 (Figure 6).

**Figure 6.**
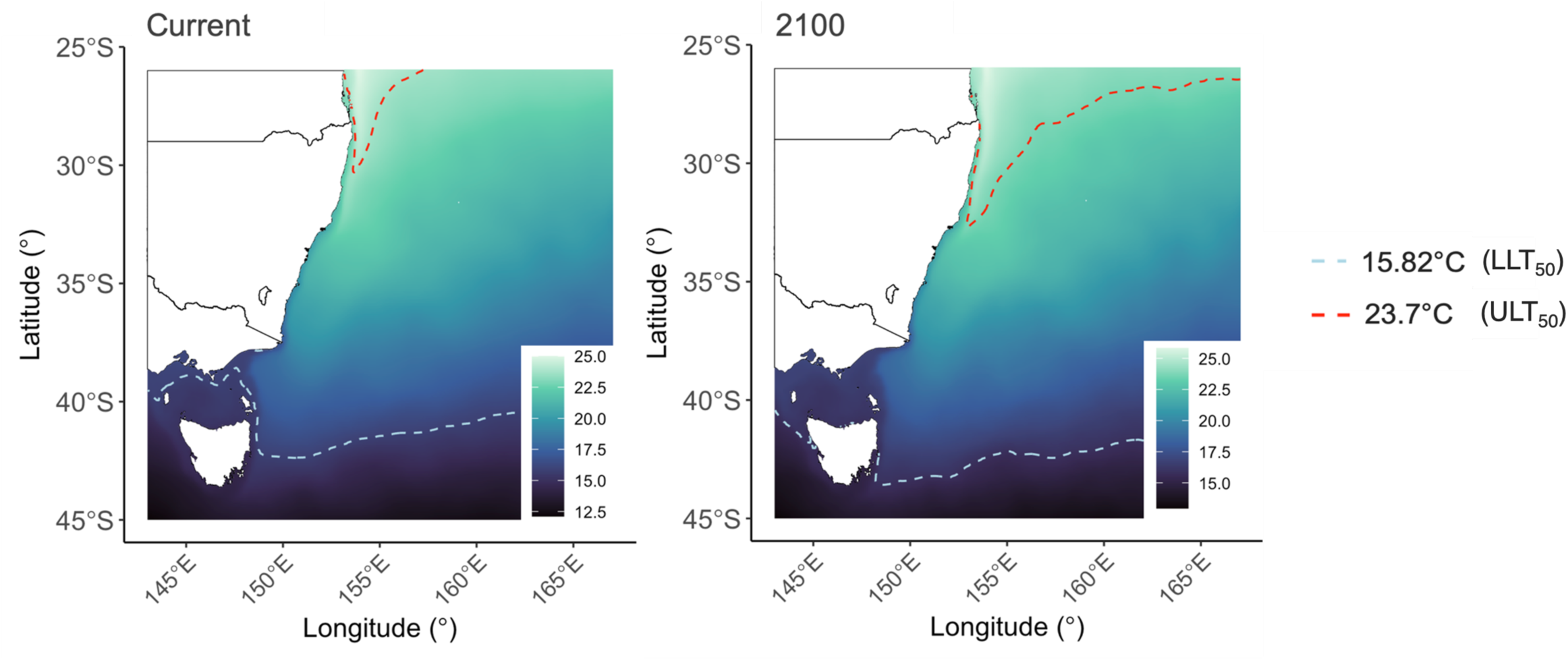
Current (left) and future (IPCC pathway SSP2-4.5) (right) dispersal potential of *Tripneustes australiae* with respect to ULT50 (23.7°C) and LLT50 (15.82°C). Background colours represent the mean SST from 2000 to 2020 extracted from Bio- Oracle SST (Assis et al. 2024).

## Discussion

As species redistribute globally, the extent and trajectories of redistributions are mediated by factors such as propagule thermal tolerance, hydrodynamic dispersal, availability of receiving habitat and successful establishment (Sunday et al. 2012; Bates et al. 2014; MacLean and Beissinger 2017; Stuart-Smith et al. 2017; Kingsbury et al. 2020; Coni et al. 2021; Byrne et al. 2022; Zarzyczny et al. 2023; Cooney et al. 2024). Here we show that the thermally suitable habitat for the early development of *T. australiae* might increase poleward in the next 100 years. For *T. australiae* there is extensive rocky reef and seagrass habitat south (poleward) of its current range. Considering the estimated planktonic duration of *T. australiae* (30 days) (Byrne et al. 2008a, b), and the optimal thermal window (17 to 23°C) it is likely that this species will shift poleward as the region warms with extension of the EAC (Sen Gupta et al. 2021; Li et al. 2022).However, our models of future dispersal potential do not account for other factors that potentially also mediate recruitment success including the presence of suitable habitat, competition with resident species and predation post-settlement (Bates et al. 2014; Coni et al. 2021; Miller et al. 2023; Zarzyczny et al. 2023). Indeed, other factors such as adaption and acclimation to future temperatures may also mediate the dispersal of *T. australiae* in the future. The optimal thermal window of *T. australiae* survival and normal development to the echinopluteus larva determined here are congruent with the realised thermal niche of this species based on the adult distribution (McLaren et al. 2025). This suggests that *T. australiae* have a similar thermal window in both the early planktonic and benthic life phases.

This study suggests that the lower thermal limit of 50% survival (LLT_50_) for *T. australiae* larvae is ∼16°C (Figure 4). In comparison to other sympatric echinoids in NSW with similar planktonic larval duration (PLD) and feeding strategies this is somewhat cold intolerant (Gall 2021; Byrne et al. 2022). For example, the lower limit of 50% survival in *C. rodgersii* is 12.6°C, while *Heliocidaris tuberculata* is 14.8°C. Both *C. rodgersii* and *H. tuberculata* currently have adult distributions similar to *T. australiae* in NSW. This includes dispersal across the Tasman onto the offshore Lord Howe and Norfolk Islands (Byrne et al. 2024; McLaren et al. 2025; Byrne et al. 2026). The southernmost observations of this species occur around Narooma where the long-term temperature range is 14.2-23.4°C (Figure 1; McLaren et al. 2025). As such, cold sea water temperatures in southeast Australia may currently limit the successful recruitment of *T. australiae* Narooma. This is interesting considering the increasingly favourable thermal conditions for the poleward larval dispersal and post larval success of *C. rodgersii* in Tasmania (Byrne et al. 2022), and seems to suggest that like *H. tuberculata*, *T. australiae* is less tolerant to the cold.

Here, we show that dispersal of *T. australiae* within their LLT_50_ is currently possible across parts of the Bass Straight, mediated by the southern limits of the EAC. However, although propagules may disperse in the EAC across the Bass Straight, the average temperatures of nearshore Tasmania are not suitable for successful development and recruitment into an adult population. The waters off southeast Australia are warming at 0.2°C per decade (CSIRO 2022) and we show that *T. australiae* may be able to successfully develop to the echinopluteus stage in the nearshore habitat in northeastern Tasmania and the islands in Bass Strait by 2100.

However, factors contributing to the successful settlement, and establishment of an adult population, will mediate the future population dynamics of *T. australiae* in southeast Australia, and should be a focus for future research. We suggest that future studies should consider the temperature limits within which juveniles recruit and survive into the benthos. This may play an important role in the boom-bust dynamics of this species, and its impacts on benthic community dynamics.

Development of *Tripneustes australiae* was tolerant to at least 3°C degrees above the parents’ ambient temperatures at the time of collection and spawning (Figure 3). Considering time of collection and spawning, *T. australiae* is unlikely to experience temperatures above 23°C in the water column in its mid to southern range. However, there is some discordance regarding the spawning periodicity of this species (O’Connor et al. 1978; Bové 2004; Lawrence and Agatsuma 2020). Some studies of *T. australiae* (then *T. gratilla*), suggest that populations in Solitary Islands Marine Park in the north of its distribution have mature gonads late summer - autumn (O’Connor et al. 1978). Other studies of populations further south in Port Stephens report mature gonads in April, while populations in Sydney begin to develop mature gonads in April through winter (Bové 2004). This suggests *T. australiae* has a prolonged spawning period, as noted for other *Tripneustes* species (Lawrence and Agatsuma 2020). As such, populations of *T. australiae* in the north in SIMP that spawn in summer may experience temperatures above 24°C (Malcolm et al. 2011, 2026; Lachs et al. 2021). The reproductive cycle of *T. australiae* in southeast Australia is an important knowledge gap to address, to understand the timing of larvae in the plankton.

Parent conditions, and where in a species’ distribution propagules originate can influence their larval thermal tolerance (Byrne 2011b; Pecorino et al. 2014; Ross et al. 2016; Byrne et al. 2019; Minuti et al. 2022; Johnson & Wong 2026). Propagules of sea urchins from different regions of their range can therefore have different developmental thermal tolerance windows (O’Connor et al. 2007; Pecorino et al. 2014). For example, propagules from parents residing at warm range edges can be more thermo-tolerant than propagules from cooler areas (Byrne 2011b). As such, in eastern Australia, due to the poleward flow of the EAC, warm edge populations can act as a source of warm-tolerant propagules (Byrne 2011b; Suthers et al. 2011; Pecorino et al. 2014). The parents used for this experiment were collected around the mid-point of this species’ distribution (McLaren et al. 2025), and so our results are likely to represent the thermal tolerance of larvae from the mid-range of this species. However, as our experiments only used two fertilizations of 2-3 males and females each, the results would not represent the variability of parental effects or genetic variability of this population. Considering the potential for different spawning periods between northern and southern populations as well as latitudinal differences in local temperature, it would be of interest to determine the thermal tolerances of larvae from range edge populations of *T. australiae* as well as incorporate more gamete sources in fertilisations to enhance representation of population level thermal tolerances. This would also allow a better understanding of cool-tolerance and range expansion potential in southern populations, and how warm-range populations may provide propagules to poleward populations.

As is typical for larval echinoderms (Byrne 2011a, b; Byrne et al. 2016, 2022; Lamare et al. 2018; Karelitz 2019; Balogh 2020; Gall 2021; Collin et al. 2021), progression through the developmental stages was faster at warmer temperatures for *T. australiae*. This has important implications for potential range extensions and local extinctions as it can affect the amount of time spent in the plankton at respective cool and warm range edges (Byrne et al. 2017; Karelitz 2019). For example, the faster a larva can develop into a pluteus, the quicker they develop long echinopluteus arms necessary for defence against predators (Vaughn and Allen 2010; Pennington et al. 1986). Propagules produced in colder waters at higher latitudes will develop more slowly and spend longer in the plankton (Byrne et al. 2017; Karelitz 2019). As such, larvae from cool-range edges may have the capacity to disperse further in the water column. We noted that *T. australiae* appears to arrest development at 14°C. However, this suggestion would need to be confirmed through longer observation of the embryos to discern between arrested development, which is fatal, and slow development. We found a 2-day developmental lag between the warm and the cold treatments. Notably, larvae in the 23°C rearing temperature were plutei by day 3, whereas larvae in the 17°C treatment didn’t reach pluteus stage until day 5. As such larvae in the water column between 16 and 17°C such as those between Sydney and Narooma during winter (Huang et al. 2017), may have the capacity to disperse further. However, it must be noted that for survival, average temperatures as we used to predict dispersal potential do not consider thermal extremes (cold or hot). Future studies should consider the effects of cold- and heat stress events on the development of *T. australiae*.

The temperature range for the successful development of *T. australiae,* (larval thermal window) narrowed through time. This suggests that cumulative thermal stress increased as larvae progressed through development, resulting in a in a narrower thermal window. This is typical of marine invertebrates, and the thermal windows of local echinoderms typically narrow through development into settlement (Przeslawski et al. 2014; Balogh et al. 2020; Gall et al. 2021). Food availability can offset thermal stress (Soars et al. 2009; Sokolova et al. 2012; Munstermann et al. 2025) and impact echinopluteus profiles (Byrne et al. 2008b). *Tripneustes australiae* have a relatively long facultative feeding period, which may prolong dispersal without feeding during early development (Byrne et al. 2008a, b). However, this study only considered larval thermal tolerances of unfed *T. australiae* larvae within its facultative feeding period (Byrne et al. 2008a, b). The lack of a food source for the larvae in our treatments differs from natural conditions, where accrued nutrient reserves likely enhance larval resilience to warming. Thus, we may have underestimated the upper thermal limits in our experiment. However, in the cooler treatments maternal reserves may not have been depleted due to slower development and so the lower thermal tolerances we present are unlikely to have been determined by food limitation.

## Conclusions

Here we established the larval thermal tolerances of *T. australiae,* which suggests that it may have a narrow thermal window in its early life history, a trait that is also reflected in its adult distribution (McLaren et al. 2025). As *T. australiae* has a narrow thermal window compared to *C. rodgersii,* it is unlikely to play a role in generation of extensive barrens-habitat poleward of its current range edge as *C. rodgersii* has done (Ling 2008). However as environmental conditions become more thermally suitable, *T. australiae* might undergo a poleward range extension in the coming decades into macroalgal-habitats that are already vulnerable to grazing by *C. rodgersii* (Glasby and Gibson 2020; Davis et al. 2023), acknowledging that successful recruitment is modulated by many factors (see Gosselin and Qian 1996; Cooney et al. 2024). As population outbreaks of *T. australiae* may have the capacity to impact already vulnerable ecosystems in southeast Australia, the mechanisms driving these outbreaks require further research.

## Funding

This research was financially supported by the Australian Government Research Training Program (ATP) PhD scholarship, Ruhm Award (University of Sydney, Marine Science Institute), the Murray Write Up Scholarship (University of Sydney) and the Holsworth Wildlife Research Endowment – Equity Trustees Charitable Foundation & the Ecological Society of Australia to EM; an Australian Research Council Discovery Early Career Research Award (DE230100141) to BS.

## Acknowledgements

The authors would like to acknowledge Matt Clements, Monique Webb and Davis Jensen for assistance in field collections.

## Conflict of interest declaration

The authors declare we have no competing interests.

## Data accessibility

The datasets supporting this article are available on the Dryad repository (http://datadryad.org/share/LINK_NOT_FOR_PUBLICATION/b6HEjIzjndWWFMCVCZY 7r8nmAdqpXqmRX7ganhZ5NUE)

## Authors’ contributions

EM: data collection, investigation, conceptualization, funding, data curation, formal analysis, methodology, visualization, writing-original draft, writing- review and editing. MB: conceptualization, data collection, supervision, writing-review and editing. PS: conceptualization, data collection. BS: supervision, writing-review and editing.

## Supplementary Material

**Figure S1.**
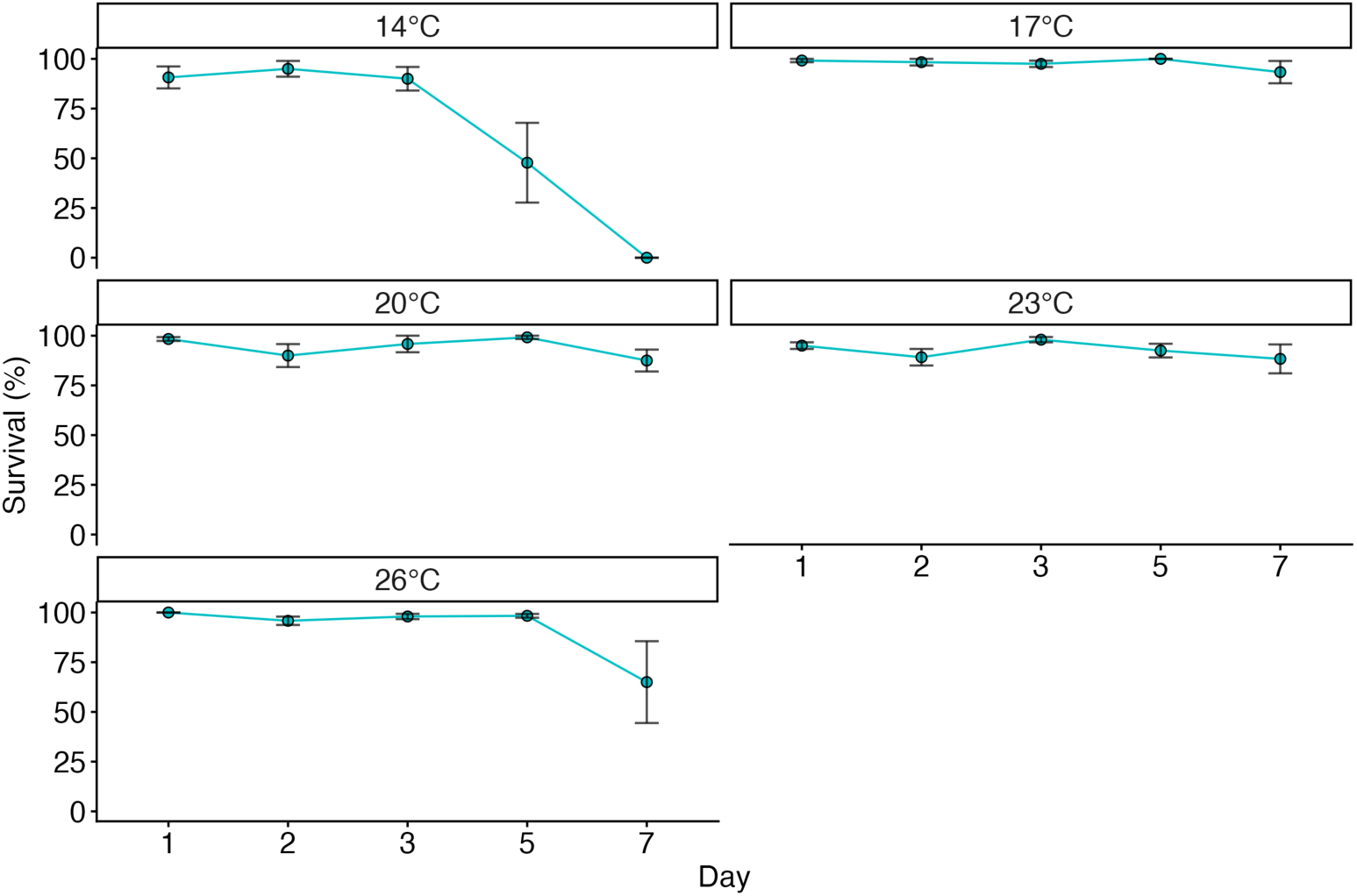
Raw data used in survival analysis. Survival (percentage alive) (±SEM) of *Tripneustes australiae* for the five time points (1-7 days post fertilisation) across five rearing temperatures (14, 17, 20, 23, & 26°C).

**Figure S2.**
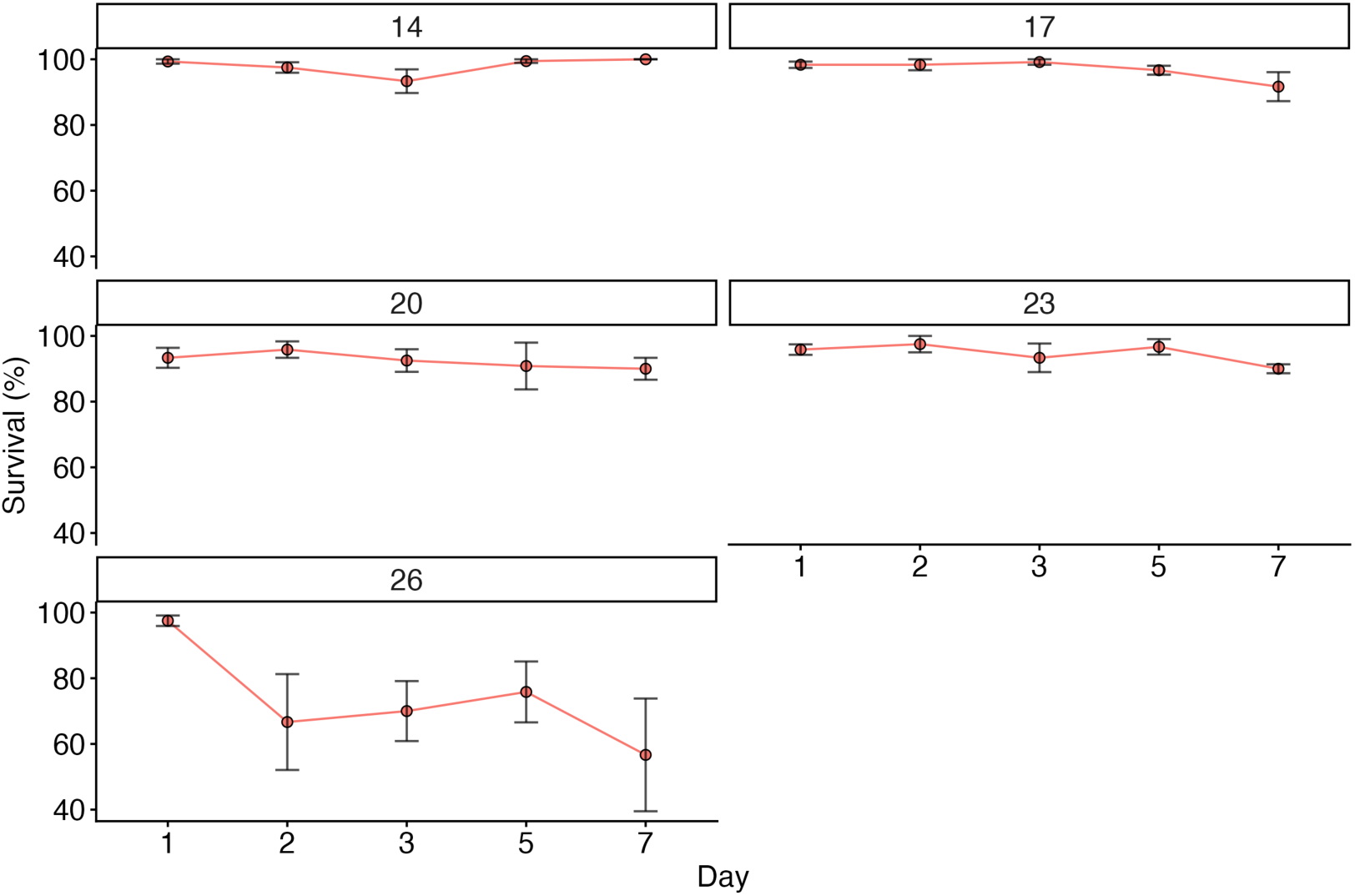
Raw data used in normality analysis. Normality (percentage normal) (±SEM) of *Tripneustes australiae* for the five time points (1-7 days post fertilisation) across five rearing temperatures (14, 17, 20, 23, & 26°C).

**Table S1.**
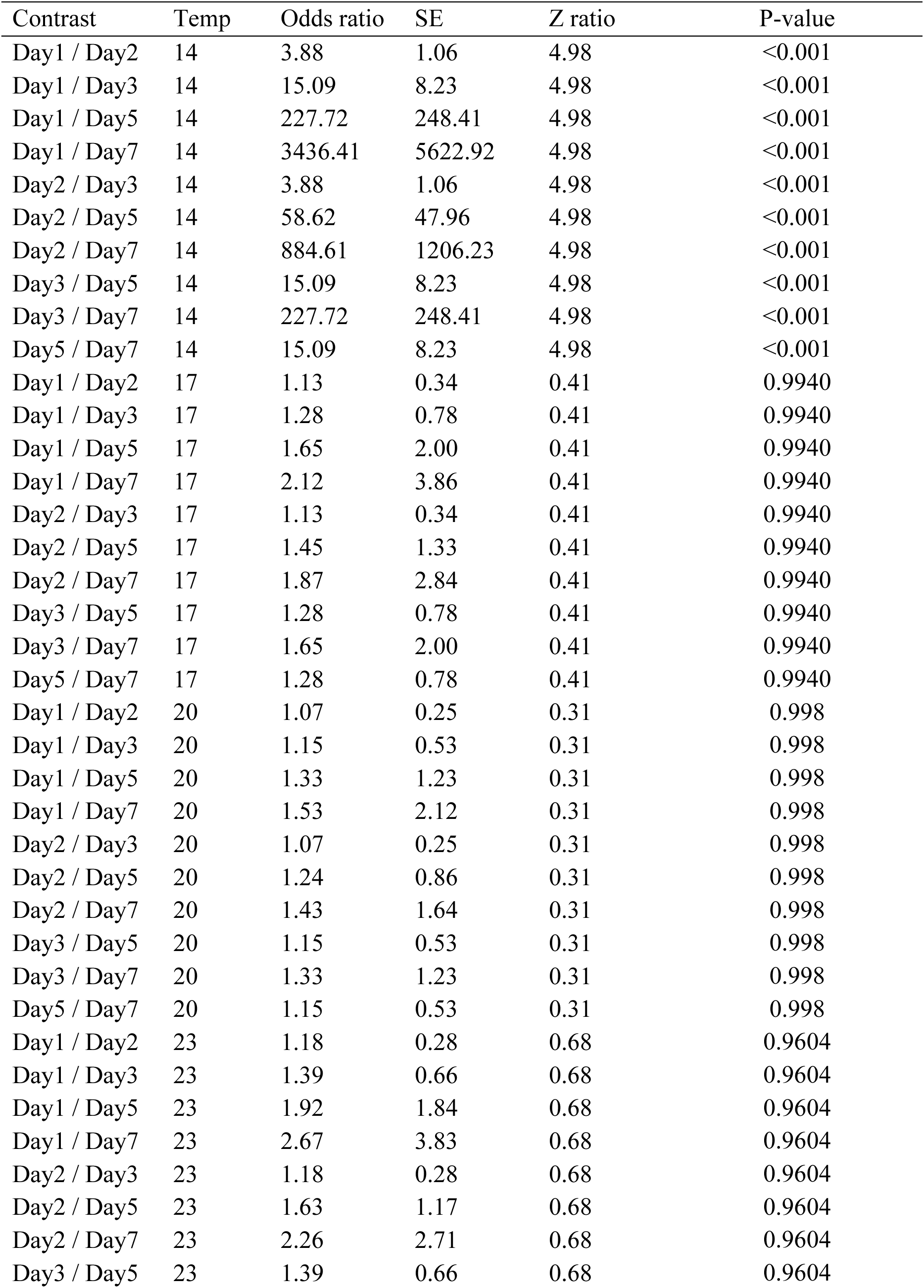
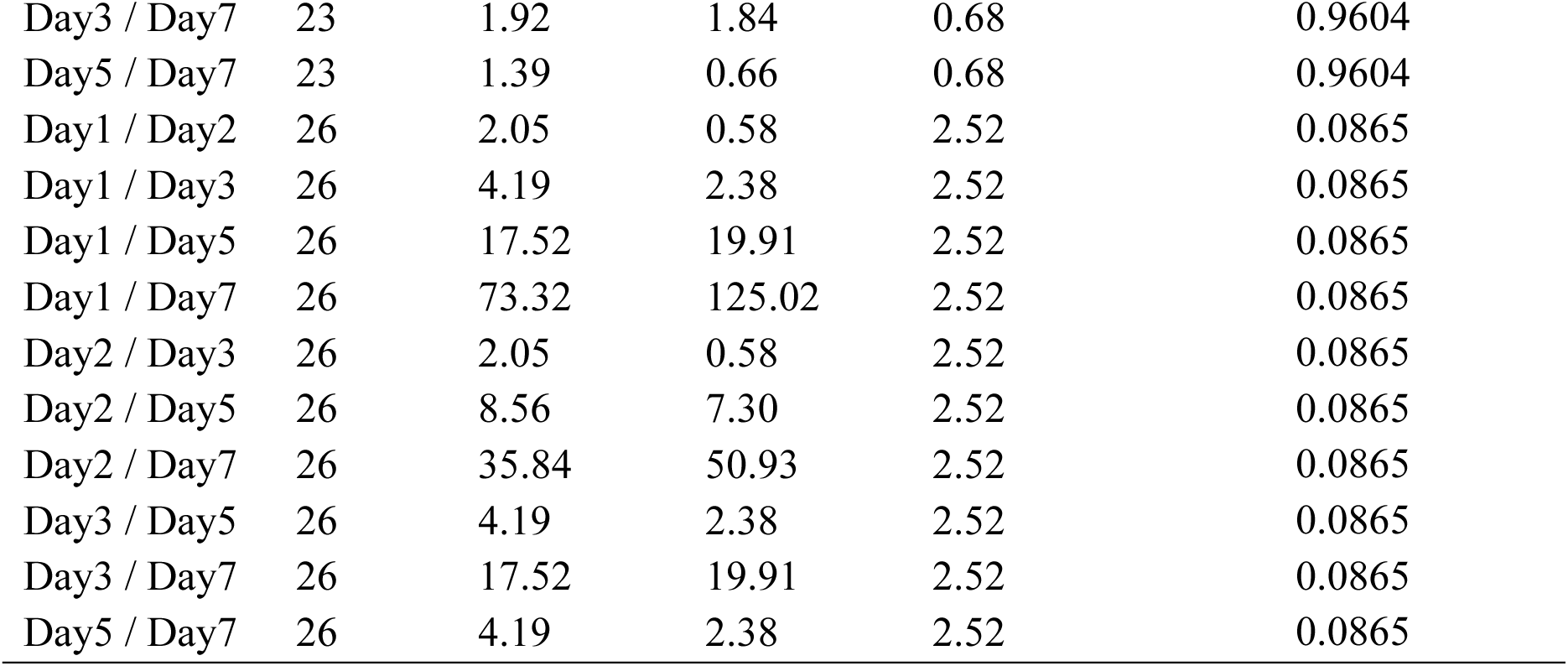
Post-hoc pairwise comparisons for survival between day post fertilisation and rearing temperatures.

**Table S2.**
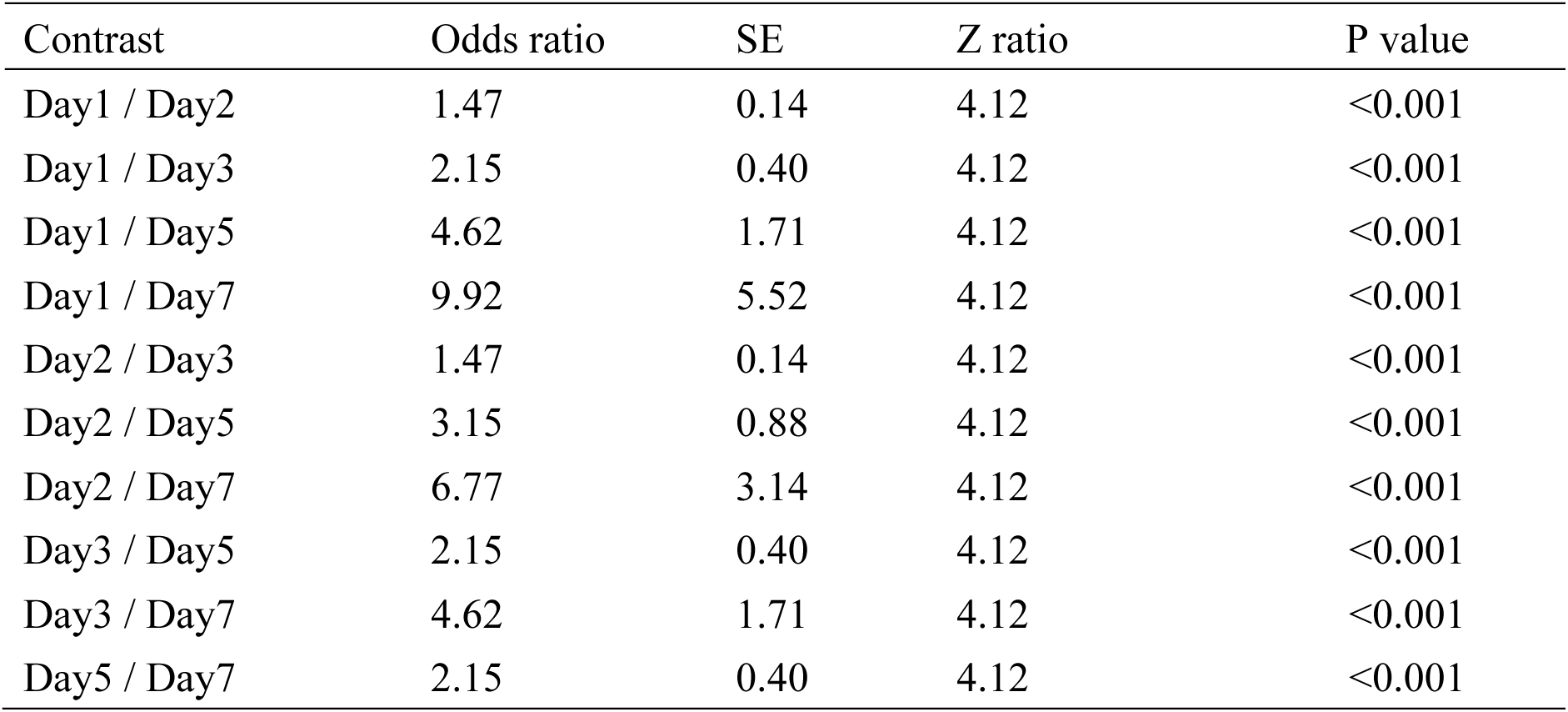
Post-hoc pairwise comparisons for normality between day post fertilisation averaged across all temperatures.

| Contrast | Odds ratio | SE | Z ratio | P value |
| --- | --- | --- | --- | --- |
| Day1 / Day2 | 1.47 | 0.14 | 4.12 | <0.001 |
| Day1 / Day3 | 2.15 | 0.40 | 4.12 | <0.001 |
| Day1 / Day5 | 4.62 | 1.71 | 4.12 | <0.001 |
| Day1 / Day7 | 9.92 | 5.52 | 4.12 | <0.001 |
| Day2 / Day3 | 1.47 | 0.14 | 4.12 | <0.001 |
| Day2 / Day5 | 3.15 | 0.88 | 4.12 | <0.001 |
| Day2 / Day7 | 6.77 | 3.14 | 4.12 | <0.001 |
| Day3 / Day5 | 2.15 | 0.40 | 4.12 | <0.001 |
| Day3 / Day7 | 4.62 | 1.71 | 4.12 | <0.001 |
| Day5 / Day7 | 2.15 | 0.40 | 4.12 | <0.001 |

**Table S3.** Post-hoc pairwise comparisons for normality between temperatures averaged across all days post fertilisation.

| Contrast | Odds ratio | SE | Z ratio | P- value |
| --- | --- | --- | --- | --- |
| Temp14 / Temp17 | 1.125315749 | 0.56810072 | 0.23386504 | 1.00 |
| Temp14 / Temp20 | 0.578264281 | 0.27485893 | -1.1523344 | 0.78 |
| Temp14 / Temp23 | 0.713528027 | 0.34048713 | -0.7073385 | 0.95 |
| Temp14 / Temp26 | 0.134884271 | 0.05920408 | -4.564192 | <0.001 |
| Temp17 / Temp20 | 0.513868469 | 0.21479356 | -1.5928197 | 0.50 |
| Temp17 / Temp23 | 0.634069173 | 0.26642973 | -1.084264 | 0.81 |
| Temp17 / Temp26 | 0.119863488 | 0.04643658 | -5.475826 | <0.001 |
| Temp20 / Temp23 | 1.233913369 | 0.47285332 | 0.54849386 | 0.98 |
| Temp20 / Temp26 | 0.233257138 | 0.0794397 | -4.2740886 | <0.001 |
| Temp23 / Temp26 | 0.189038504 | 0.06551976 | -4.8062018 | <0.001 |

**Table S4.**
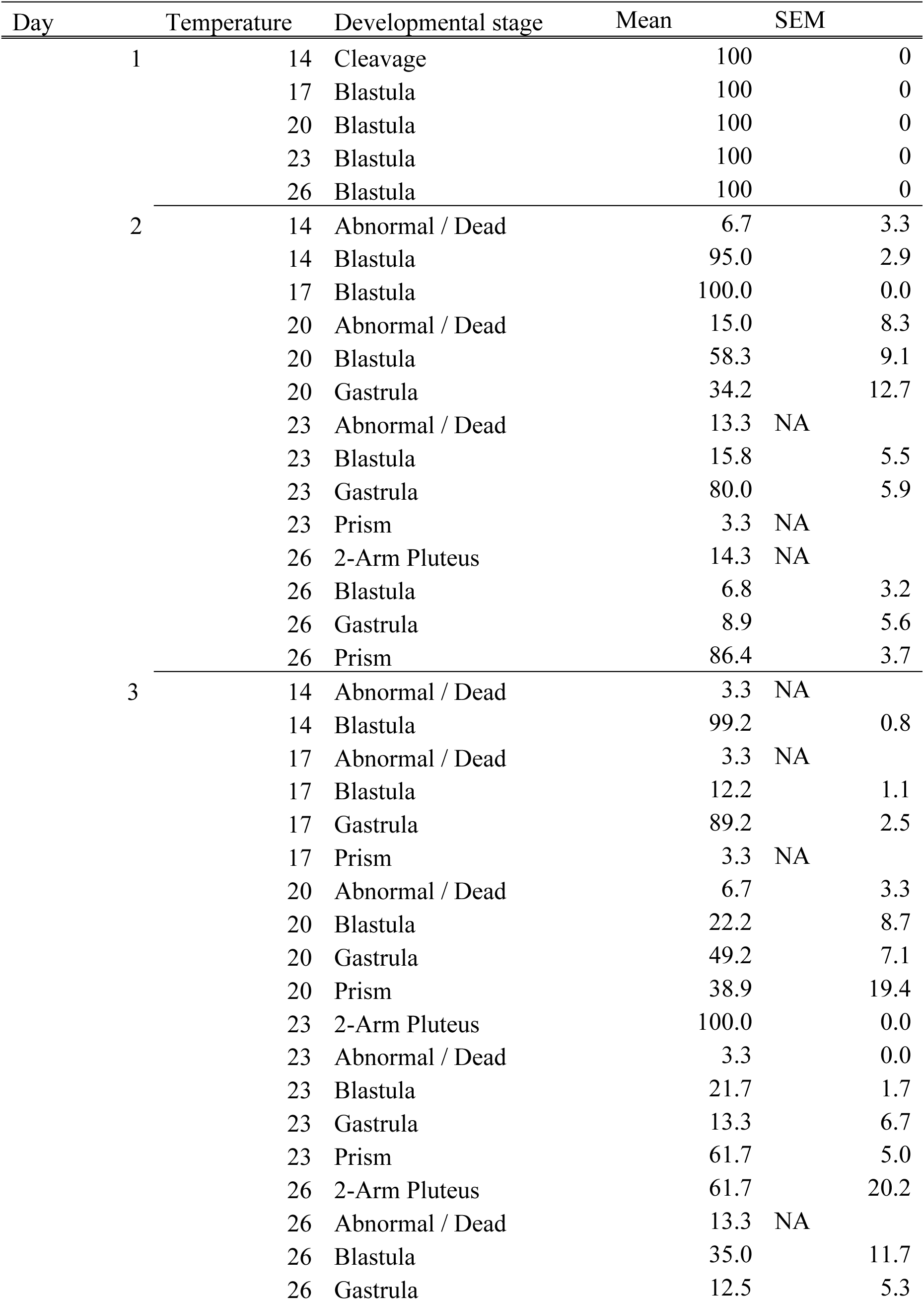
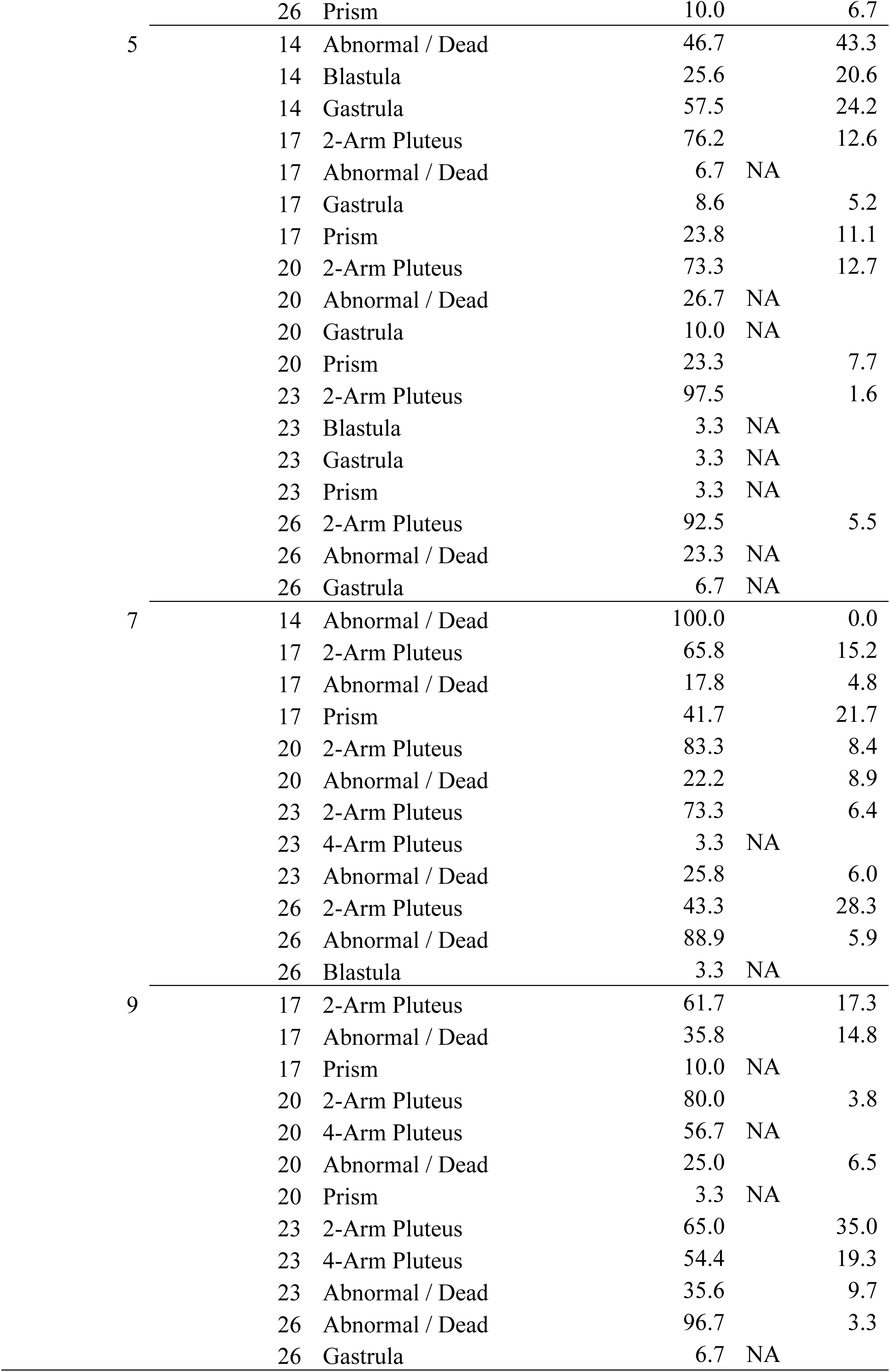
The mean number of *Tripneustes australiae* larvae across replicates (± SEM, n = 4) at different developmental stages for each day post-fertilization (n = 6) and rearing temperature (n = 5).

## Notes

### Competing Interest Statement

The authors have declared no competing interest.

## References

1. Assis J, Fernández Bejarano SJ, Salazar VW, Schepers L, Gouvêa L, Fragkopoulou E, Leclercq F, Vanhoorne B, Tyberghein L, Serrão EA, Verbruggen H, De Clerck O (2024) Bio-ORACLE v3.0. Pushing marine data layers to the CMIP6 Earth System Models of climate change research. Global Ecol Biogeogr 33:e13813. doi: 10.1111/geb.13813

2. Balogh R, Byrne B (2020) Developing in a warming intertidal, negative carry over effects of heatwave conditions in development to the pentameral starfish in *Parvulastra exigua*. Mar Env Res 162: 105083. doi: 10.1016/j.marenvres.2020.105083

3. Bates AE, Pecl GT, Frusher S, Hobday AJ, Wernberg T, Smale DA, Sunday JM, Hill NA, Dulvy NK, Colwell RK, Holbrook NJ, Fulton EA, Slawinski D, Feng M, Edgar GJ, Radford BT, Thompson PA, Watson RA (2014) Defining and observing stages of climate-mediated range shifts in marine systems. Glob Environ Change 26:27–38. doi: 10.1016/j.gloenvcha.2014.03.009

4. Beachwatch (NSW) (2024) Water quality data. New South Wales Government. Retrieved [2024]. https://www.beachwatch.nsw.gov.au/waterMonitoring/waterQualityData

5. Bové U (2004) Annual reproductive cycle of Tripneustes gratilla (Linneus, 1758) and the effect of temperature on sperm concentration, early cleavage and larval development. Honours Thesis. Department of Biological Sciences, Macquarie University, NSW 2109

6. Bronstein O, Kroh A, Tautscher B, Liggins L, Haring E (2017) Cryptic speciation in pan- tropical sea urchins: a case study of an edge-of-range population of *Tripneustes* from the Kermadec Islands. Sci Rep 7:5948. doi: 10.1038/s41598-017-06183-2

7. Bronstein O, Kroh A, Miskelly AD, Smith SDA, Dworjanyn SA, Mos B, Byrne M (2019) Implications of range overlap in the commercially important pan-tropical sea urchin genus *Tripneustes* (Echinoidea: Toxopneustidae). Mar Biol 166:34. doi: 10.1007/s00227-019-3478-4

8. Brooks M E, Kristensen K, Benthem K J, van, Magnusson A, Berg C W, Nielsen A, Skaug H J, Mächler M, Bolker B M (2017) glmmTMB Balances Speed and Flexibility Among Packages for Zero-inflated Generalized Linear Mixed Modeling. R J 9:378. doi: 10.32614/RJ-2017-066

9. Byrne M (2011a) Impact of ocean warming and ocean acidification on marine invertebrate life history sages: Vulnerabilities and potential for persistence in a changing ocean. Oceanography Mar Biol: Annual Rev 49: 1–42. R. Eds: N. Gibson, R. J. A. Atkinson. Taylor & Francis.

10. Byrne M (2011b) Sea urchin development in a global change hotspot, potential for southerly migration of thermotolerant propagules. Deep Sea Res II 58: 712–719. doi: 10.1016/j.dsr2.2010.06.010

11. Byrne M, O’Hara T (eds) (2017) Australian Echinoderms: Biology, Ecology and Evolution. CSIRO Publishing, Clayton South VIC 3169 Australia

12. Byrne M, Przeslawski R (2013) Multistressor impacts of warming and acidification of the ocean on marine invertebrates’ life histories. Int Comp Biol 53 (4): 582–596 doi:10.1093/icb/ict049

13. Byrne M, Prowse TAA, Sewell MA, Dworjanyn S, Williamson JE, Vaïtilingon D (2008a) Maternal provisioning for larvae and larval provisioning for juveniles in the toxopneustid sea urchin *Tripneustes gratilla*. Mar Biol 155:473–482. doi: 10.1007/s00227-008-1045-5

14. Byrne M, Sewell MA, Prowse TAA (2008b) Nutritional ecology of sea urchin larvae: influence of endogenous and exogenous nutrition on echinopluteal growth and phenotypic plasticity in *Tripneustes gratilla*. Funct Ecol 22:643–648. doi: 10.1111/j.1365-2435.2008.01427.x

15. Byrne M, Gall ML, Wolfe K, Agüera A (2016) From pole to pole: the potential for the Arctic seastar *Asterias amurensis* to invade a warming Southern Ocean. Glob Change Biol (2016) 22, 3874–3887, doi: 10.1111/gcb.13304

16. Byrne M, Ross PM, Dworjanyn SA, Parker L (2017) Larval ecology in the face of changing climate—Impacts of ocean warming and ocean acidification. In: Evolutionary Ecology of Marine Invertebrate Larvae. Oxford University Press, pp.251–272

17. Byrne M, Gall ML, Campbell H, Lamare MD, Holmes SP (2022) Staying in place and moving in space: Contrasting larval thermal sensitivity explains distributional changes of sympatric sea urchin species to habitat warming. Glob Change Biol 28:3040–3053. doi: 10.1111/gcb.16116

18. Byrne M, McLaren EJ, Wilson L, Webb M, Eichman S, Fraser N, Lenehan N, Miskelly A, Shaw I, Turnbull J, Sommer B (2024) Population density fluctuations of the sea urchin *Tripneustes australiae* in New South Wales, Australia. Cah. Biol. Mar. (2024) 65: 377–388 doi: 10.21411/CBM.A.43CEC0F7

19. Byrne M, McLaren E, Wilson L, Cooper A, Davey M, Sommer B (2026) Return of the lamingtons–boom-and-bust populations of the barrens forming sea urchin *Tripneustes australiae* in Lord Howe Island Marine Park. Coral Reefs. doi: 10.1007/s00338-026-02851-w

20. Castro LC, Cetina-Heredia P, Roughan M, Dworjanyn S, Thibaut L, Chamberlain MA, Feng M, Vergés A (2020) Combined mechanistic modelling predicts changes in species distribution and increased co-occurrence of a tropical urchin herbivore and a habitat- forming temperate kelp. Divers Distrib 26:1211–1226. doi: 10.1111/ddi.13073

21. Collin R, Rebolledo AP, Smith E, Chan KYK (2021) Thermal tolerance of early development predicts the realized thermal niche in marine ectotherms. Funct Ecol 35:1679–1692. doi: 10.1111/1365-2435.13850

22. Coni EOC, Booth DJ, Nagelkerken I (2021) Novel species interactions and environmental conditions reduce foraging competency at the temperate range edge of a range- extending coral reef fish. Coral Reefs 40:1525–1536. doi: 10.1007/s00338-021-02150-6

23. Cooney C, Sommer B, Marzinelli EM, Figueira WF (2024) The role of microbial biofilms in range shifts of marine habitat-forming organisms. Trends Microbiol 32:190–199. doi: 10.1016/j.tim.2023.07.015

24. CSIRO & Bureau of Meterology (2022) State of The Climate 2022. (Report No. 22-00220_OA). https://www.bom.gov.au/climate

25. Davis TR, Knott NA, Champion C, Przeslawski R (2023) Impacts of climate change on densities of the urchin *Centrostephanus rodgersii* vary among marine regions in eastern Australia. Diversity 15:419. doi: 10.3390/d15030419

26. Edgar GJ, Stuart-Smith RD, Heather FJ, Barrett NS, Turak E, Sweatman H, Emslie MJ, Brock DJ, Hicks J, French B, Baker SC, Howe SA, Jordan A, Knott NA, Mooney P, Cooper AT, Oh ES, Soler GA, Mellin C, Ling SD, Dunic JC, Turnbull JW, Day PB, Larkin MF, Seroussi Y, Stuart-Smith J, Clausius E, Davis TR, Shields J, Shields D, Johnson OJ, Fuchs YH, Denis-Roy L, Jones T, Bates AE (2023) Continent-wide declines in shallow reef life over a decade of ocean warming. Nature 615:858–865. doi: 10.1038/s41586-023-05833-y

27. Fox J, Weisberg S (2019) An R Companion to Applied Regression, Third edition. Sage, Thousand Oaks CA.

28. Fredston A, Tingley MW, Neate-Clegg MH, Evans LJ, Antão LH, Ban NC and Scheffers, BR (2025) Reimagining species on the move across space and time. TREE 40 (7): 629– 641. doi: 10.1016/j.tree.2025.03.015

29. Gall ML (2021) Effects of marine heatwave conditions across the metamorphic transition to the juvenile sea urchin (*Heliocidaris erythrogramma*). Mar Pol Bul 163: 111914. doi: 10.1016/j.marpolbul.2020.111914

30. García Molinos J, Halpern BS, Schoeman DS, Brown CJ, Kiessling W, Moore PJ, Pandolfi JM, Poloczanska ES, Richardson AJ, Burrows MT (2016) Climate velocity and the future global redistribution of marine biodiversity. Nature Clim Change 6:83–88. doi: 10.1038/nclimate2769

31. Glasby TM, Gibson PT (2020) Decadal dynamics of subtidal barrens habitat. Mar Environ Res 154:104869. doi: 10.1016/j.marenvres.2019.104869

32. Gosselin LA, Qian PY (1996) Early post-settlement mortality of an intertidal barnacle: a critical period for survival. MEPS. 135:69–75. doi:10.3354/mwpa135069

33. Hartig F (2026) DHARMa: Residual diagnostics for hierarchical (multi-Level / mixed) regression models. v 0.4.7.

34. Hobday AJ, Pecl GT (2014) Identification of global marine hotspots: sentinels for change and vanguards for adaptation action. Rev Fish Biol Fisheries 24:415–425. doi: 10.1007/s11160-013-9326-6

35. Huang B, Thorne PW, Banzon VF, Boyer T, Chepurin G, Lawrimore JH, Menne MJ, Smith TM, Vose RS, Zhang H-M (2017) Extended reconstructed sea surface temperature, Version 5 (ERSSTv5): Upgrades, validations, and intercomparisons. J Clim 30:8179–8205. doi: 10.1175/JCLI-D-16-0836.1

36. IPCC (2023) Climate Change 2021 – The physical science basis: Working group I contribution to the sixth assessment report of the Intergovernmental Panel on Climate Change, 1st edn. Cambridge University Press

37. Jones GP, Andrew NL (1990) Herbivory and patch dynamics on rocky reefs in temperate Australasia: The roles of fish and sea urchins. Austral Ecol 15:505–520. doi: 10.1111/j.1442-9993.1990.tb01474.x

38. Johnson EC, Wong JM (2026) From stress to response: a systematic review of epigenetic pathways underlying gene expression and phenotypic plasticity in aquatic invertebrates. Front Mar Sci 13:1767697. doi: 10.3389/fmars.2026.1767697

39. Karelitz S, Lamare M, Mos B, De Bari H, Dworjanyn SA, Byrne M(2019) Impact of growing up in a warmer, lower pH future on offspring performance: transgenerational plasticity in a pan-tropical sea urchin. Coral Reefs 38:1085–1095. doi: 10.1007/s00338-019-01855-z

40. Kingsbury KM, Gillanders BM, Booth DJ, Nagelkerken I (2020) Trophic niche segregation allows range-extending coral reef fishes to co-exist with temperate species under climate change. Glob Change Biol 26:721–733. doi: 10.1111/gcb.14898

41. Lachs L, Sommer B, Cant J, Hodge JM, Malcolm HA, Pandolfi JM, Beger M (2021) Linking population size structure, heat stress and bleaching responses in a subtropical endemic coral. Coral Reefs 40:777–790. doi: 10.1007/s00338-021-02081-2

42. Lamare M, Harianto J, Uthicke S, Agüera A, Karelitz S, Pecorino D, Chin J, Byrne M (2018) Larval thermal windows in native and hybrid *Pseudoboletia* progeny (Echinoidea) as potential drivers of the hybridization zone. Mar Ecol Prog Ser 598:99–112. doi: 10.3354/meps12601

43. Lang BJ, Donelson JM, Bairos KR, Wheeler CR, Caballes CF, Uthicke S, Pratchett MS (2023) Impacts of ocean warming on echinoderms: A meta-analysis. Ecol Evol 13:e10307. doi: 10.1002/ece3.10307

44. Lawrence JM, Agatsuma Y (2020) Tripneustes. In: Sea Urchins: Biology and Ecology. Elsevier, pp 681–703.

45. Lenth R (2024) emmeans: Estimated Marginal Means, aka Least-Squares Means. R package version 1.9.0.

46. Li J, Roughan M, Kerry C (2022) Drivers of ocean warming in the western boundary currents of the Southern Hemisphere. Nat Clim Chang 12:901–909. doi: 10.1038/s41558-022-01473-8

47. Ling SD (2008) Range expansion of a habitat-modifying species leads to loss of taxonomic diversity: a new and impoverished reef state. Oecologia 156:883–894. doi: 10.1007/s00442-008-1043-9

48. MacLean SA, Beissinger SR (2017) Species’ traits as predictors of range shifts under contemporary climate change: A review and meta-analysis. Glob Change Biol23:4094– 4105. doi: 10.1111/gcb.13736

49. Malcolm HA, Davies PL, Jordan A, Smith SDA (2011) Variation in sea temperature and the East Australian Current in the Solitary Islands region between 2001–2008. Deep-Sea Res 58:616–627. doi: 10.1016/j.dsr2.2010.09.030

50. Malcolm HA, Lechene M, Ferrari R, Champion C, Dalton SJ, Beger M, Cant J, Chong F, Hsiao WV, Kim SW, Lachs L, Smith SDA, Figueira WF, Sommer B (2026) Multi-decadal resilience (2002–2023) of subtropical coral communities in the Solitary Islands, eastern Australia, despite severe cumulative disturbances. Coral Reefs. doi: 10.1007/s00338-026-02867-2

51. McEdward LR, Miner BG (2001) Larval and life-cycle patterns in echinoderms. Can J Zool 79: 1125–1170. doi: 10.1139/cjz-79-7-1125

52. McLaren E, Bronstein O, Kroh A, Winkler V, Miskelly A, Sommer B, Byrne M (2023) Hidden in plain sight: *Tripneustes kermadecensis* (Echinodermata: Echinoidea) is a junior synonym of the eastern Australian sea urchin *Evechinus australiae* described in 1878. Invertebr Syst 37:741–754. doi: 10.1071/IS23038

53. McLaren E, Sommer B, Pandolfi JM, Beger M, Byrne M (2024) Taxa-dependent temporal trends in the abundance and size of sea urchins in subtropical eastern Australia. Ecol Evol 14:e11412. doi: 10.1002/ece3.11412

54. McLaren E, Sommer B, Pine C, Miskelly A, Byrne M (2025) Searchin’ for urchins: Utilising museum collections and citizen science to assess species on the move in the genus *Tripneustes*. J Biogeo 52:e15092. doi: 10.1111/jbi.15092

55. McLaren E, Byrne M, Pandolfi JM, Berger M, Sommer B (2026) Spatial and temporal patterns in echinoderm species and trait composition in the subtropical-to-temperate transition zone of eastern Australia. Proc R Soc B 293: 20252671. doi:10.1098/rspb.2025.2671

56. Miller MGR, Reimer JD, Sommer B, Cook KM, Pandolfi JM, Obuchi M, Beger M (2023) Temperate functional niche availability not resident-invader competition shapes tropicalisation in reef fishes. Nat Commun 14:2181. doi: 10.1038/s41467-023-37550-5

57. Minuti JJ, Byrne M, Campbell H, Hemraj DA, Russell BD (2022) Live-fast-die-young: Carryover effects of heatwave-exposed adult urchins on the development of the next generation. Glob Change Biol 28:5781–5792. doi: 10.1111/gcb.16339

58. Moreira-Saporiti A, Hoeijmakers D, Reuter H, Msuya FE, Gese K, Teichberg M (2023) Bottom-up and top-down control of seagrass overgrazing by the sea urchin *Tripneustes gratilla*. Mar Ecol. doi: 10.1111/maec.12734

59. Munstermann MJ, Karelitz SE, Ferraro R, Rogers-Bennett L, Simons RD, Okamoto DK (2026) Food limitation erodes the thermal tolerance of larvae in an ecologically influential marine herbivore. Ecology. 107(1):e70288. doi: 10.1002/ecy.70288.

60. O’Connor C, Riley G, Lefebvre S, Bloom D (1978) Environmental influences on histological changes in the reproductive cycle of four New South Wales sea urchins. Aquaculture 15:1–17. doi: 10.1016/0044-8486(78)90068-6

61. O’Connor MI, Bruno JF, Gaines SD, Halpern BS, Lester SE, Kinlan BP, Weiss JM (2007) Temperature control of larval dispersal and the implications for marine ecology, evolution, and conservation. Proc Natl Acad Sci USA 104:1266–1271. doi: 10.1073/pnas.0603422104

62. B.F. Oliveira, R. Bertrand, M.L. Pinsky, N. Casajus, B.W. Wolfe, B.R. Scheffers, F. Villalobos, G. Grenouillet, G.T. Pecl, I. Chen, J.A. Baecher, J.A. Lawlor, J. Sunday, J. Murienne, J. Rolland, L.M. Thompson, L.T. Lancaster, M.A. Rubenstein, N.A. Moore, R.M.W.J. Bandara, S.E. Diamond, S.R. Weiskopf, J. Lenoir, & L. Comte, Species range shifts often speed ahead of their modeled climatic niches, Proc. Natl. Acad. Sci. U.S.A. 123 (14) e2515903123, 10.1073/pnas.2515903123 (2026).

63. Pecorino D, Barker MF, Dworjanyn SA, Byrne M, Lamare MD (2014) Impacts of near future sea surface pH and temperature conditions on fertilisation and embryonic development in *Centrostephanus rodgersii* from northern New Zealand and northern New South Wales, Australia. Mar Biol 161:101–110. doi: 10.1007/s00227-013-2318-1

64. Pennington T, Chia F-S, Rumrill S (1986) Stage-specific predation upon embryos and larvae of the pacific sand dollar *Dendraster excentricus* by 11 species of common zooplankton predators. Bul Mar Sci 39(2): 234–240.

65. Pinsky ML, Worm B, Fogarty MJ, Sarmiento JL, Levin SA (2013) Marine taxa track local climate velocities. Science 341:1239–1242. doi: 10.1126/science.1239352

66. Poloczanska ES, Burrows MT, Brown CJ, García Molinos J, Halpern BS, Hoegh-Guldberg O, Kappel CV, Moore PJ, Richardson AJ, Schoeman DS, Sydeman WJ (2016) Responses of marine organisms to climate change across oceans. Front Mar Sci. doi: 10.3389/fmars.2016.00062

67. Przeslawski R, Chick R, Day J, Glasby T, Knott N (2023) Research Summary New South Wales Barrens. NSW Department of Primary Industries.

68. R Core Team (2025) R: A Language and Environment for Statistical Computing. Austria, Vienna.

69. Ross PM, Parker L, Byrne M (2016) Transgenerational responses of molluscs and echinoderms to changing ocean conditions. ICES Journal of Marine Science 73:537–549. doi: 10.1093/icesjms/fsv254

70. Sandblom E, Clark TD, Gräns A, Ekström A, Brijs J, Sundström LF, Odelström A, Adill A, Aho T, Jutfelt F (2016) Physiological constraints to climate warming in fish follow principles of plastic floors and concrete ceilings. Nat Commun 7:11447. doi: 10.1038/ncomms11447

71. Sen Gupta A, Stellema A, Pontes GM, Taschetto AS, Vergés A, Rossi V (2021) Future changes to the upper ocean Western Boundary Currents across two generations of climate models. Sci Rep 11:9538. doi: 10.1038/s41598-021-88934-w

72. Soars N, Prowse T, Byrne M (2009) Overview of phenotypic plasticity in echinoid larvae, ‘Echinopluteus transversus’ type vs. typical echinoplutei. Mar Ecol Prog Ser 383:113–125. doi: 10.3354/meps07848

73. Sokolova IM, Frederich M, Bagwe R, Lannig G, Sukhotin AA (2012) Energy homeostasis as an integrative tool for assessing limits of environmental stress tolerance in aquatic invertebrates. Mar Env Res 79:1–15. doi: 10.1016/j.marenvres.2012.04.003

74. Steneck RS (2020) Regular sea urchins as drivers of shallow benthic marine community structure. In: Developments in Aquaculture and Fisheries Science. Elsevier, pp 255– 279

75. Strathmann RR (1993) Hypotheses on the origins of marine larvae. Annu Rev Ecol Syst 1993. 24:89–117. doi: 0066-4162/9311120-0089

76. Stuart-Smith RD, Edgar GJ, Bates AE (2017) Thermal limits to the geographic distributions of shallow-water marine species. Nat Ecol Evol 1:1846–1852. doi: 10.1038/s41559-017-0353-x

77. Stuart-Smith RD, Edgar GJ, Clausius E, Oh ES, Barrett NS, Emslie MJ, Bates AE, Bax N, Brock D, Cooper A, Davis TR, Day PB, Dunic JC, Green A, Hasweera N, Hicks J, Holmes TH, Jones B, Jordan A, Knott N, Larkin MF, Ling SD, Mooney P, Pocklington JB, Seroussi Y, Shaw I, Shields D, Smith M, Soler GA, Stuart-Smith J, Turak E, Turnbull JW, Mellin C (2022) Tracking widespread climate-driven change on temperate and tropical reefs. Curr Biol 32:4128–4138.e3. doi: 10.1016/j.cub.2022.07.067

78. Sunday JM, Bates AE, Dulvy NK (2011) Global analysis of thermal tolerance and latitude in ectotherms. Proc R Soc B 278:1823–1830. doi: 10.1098/rspb.2010.1295

79. Sunday JM, Bates AE, Dulvy NK (2012) Thermal tolerance and the global redistribution of animals. Nature Clim Change 2:686–690. doi: 10.1038/nclimate1539

80. Sunday JM, Pecl GT, Frusher S, Hobday AJ, Hill N, Holbrook NJ, Edgar GJ, Stuart-Smith R, Barrett N, Wernberg T, Watson RA, Smale DA, Fulton EA, Slawinski D, Feng M, Radford BT, Thompson PA, Bates AE (2015) Species traits and climate velocity explain geographic range shifts in an ocean-warming hotspot. Ecol Lett 18:944–953. doi: 10.1111/ele.12474

81. Suthers IM, Young JW, Baird ME, Roughan M, Everett JD, Brassington GB, Byrne M, Condie SA, Hartog JR, Hassler CS, Hobday AJ, Holbrook NJ, Malcolm HA, Oke PR, Thompson PA, Ridgway K (2011) The strengthening East Australian Current, its eddies and biological effects — an introduction and overview. Deep-Sea Res 58:538–546. doi: 10.1016/j.dsr2.2010.09.029

82. Tuckett CA, De Bettignies T, Fromont J, Wernberg T (2017) Expansion of corals on temperate reefs: direct and indirect effects of marine heatwaves. Coral Reefs 36:947–956. doi: 10.1007/s00338-017-1586-5

83. Underwood AJ, Kingsford MJ, Andrew NL (1991) Patterns in shallow subtidal marine assemblages along the coast of New South Wales. Austral Ecol 16:231–249. doi: 10.1111/j.1442-9993.1991.tb01050.x

84. Uthicke S, Schaffelke B, Byrne M (2009) A boom–bust phylum? Ecological and evolutionary consequences of density variations in echinoderms. Ecol Monogr 79:22.

85. Valentine JP, Edgar GJ (2010) Impacts of a population outbreak of the urchin *Tripneustes gratilla* amongst Lord Howe Island coral communities. Coral Reefs 29:399–410. doi:10.1007/s00338-010-0610-9

86. Vaughn D, Allen JD (2010) The Peril of the Plankton. Int Comp Biol50:552–570. doi: 10.1093/icb/icq037

87. Vergés A, Steinberg PD, Hay ME, Poore AGB, Campbell AH, Ballesteros E, Heck KL, Booth DJ, Coleman MA, Feary DA, Figueira W, Langlois T, Marzinelli EM, Mizerek T, Mumby PJ, Nakamura Y, Roughan M, van Sebille E, Gupta AS, Smale DA, Tomas F, Wernberg T, Wilson SK (2014) The tropicalization of temperate marine ecosystems: climate-mediated changes in herbivory and community phase shifts. Proc R Soc B 281:20140846. doi: 10.1098/rspb.2014.0846

88. Vergés A, McCosker E, Mayer-Pinto M, Coleman MA, Wernberg T, Ainsworth T, Steinberg PD (2019) Tropicalisation of temperate reefs: Implications for ecosystem functions and management actions. Funct Ecol 33:1000–1013. doi: 10.1111/1365-2435.13310

89. Wang Y, Naumann U, Eddelbuettel D, Wilshire J, Warton D (2026) mvabund: Statistical Methods for Analysing Multivariate Abundance Data. v 4.2.8.

90. Wood SN (2017) Generalized additive models: an introduction with R, Second edition. CRC Press/Taylor & Francis Group, Boca Raton

91. Warton DI (2005) Many zeros does not mean zero inflation: comparing the goodness-of-fit of parametric models to multivariate abundance data. Environmetrics16: 275–289. doi:10.1002/env.702

92. Zarzyczny KM, Rius M, Williams ST, Fenberg PB (2023) The ecological and evolutionary consequences of tropicalisation. Trends Ecol Evol S0169534723002732. doi: 10.1016/j.tree.2023.10.006

